# An amino-terminal threonine/serine motif is necessary for activity of the Crp/Fnr homolog, MrpC, and for *Myxococcus xanthus* developmental robustness

**DOI:** 10.1101/737593

**Authors:** Brooke E. Feeley, Vidhi Bhardwaj, Patrick T. McLaughlin, Stephen Diggs, Gregor M. Blaha, Penelope I. Higgs

**Affiliations:** Department of Biological Sciences, Wayne State University, Detroit, MI, USA; Department of Ecophysiology, Max Planck Institute for Terrestrial Microbiology, Marburg, Hesse, Germany; Department of Biochemistry, University of California, Riverside, Riverside, CA, USA

**Keywords:** *Myxococcus xanthus*, MrpC, Crp/Fnr, development, biofilm, developmental buffer

## Abstract

The Crp/Fnr family of transcriptional regulators play central roles in transcriptional control of diverse physiological responses. Activation of individual family members is controlled by a surprising diversity of mechanisms tuned to the particular physiological responses or lifestyles that they regulate. MrpC is a Crp/Fnr homolog that plays an essential role in controlling the *Myxococcus xanthus* developmental program. A long-standing model proposed that MrpC activity is controlled by the Pkn8/Pkn14 serine/threonine kinase cascade which phosphorylates MrpC on threonine residue(s) located in its extreme amino terminus. In this study, we demonstrate that a stretch of consecutive threonine and serine residues, T_21_ T_22_ S_23_ S_24,_ is necessary for MrpC activity by promoting efficient DNA binding. Mass spectrometry analysis indicated the TTSS motif is not directly phosphorylated by Pkn14 *in vitro* but is necessary for efficient Pkn14-dependent phosphorylation on several residues in the remainder of the protein. Pkn8 and Pkn14 kinase activities do not play obvious roles in controlling MrpC activity in wild type *M. xanthus* under laboratory conditions, but likely modulate MrpC DNA binding in response to unknown environmental conditions. Interestingly, mutational analysis of the TTSS motif caused non-robust developmental phenotypes, revealing that MrpC plays a role in developmental buffering.

## INTRODUCTION

Crp/Fnr transcriptional regulators belong to a large family characterized by an amino terminal cyclic nucleotide binding (“cNMP”) domain followed by a characteristic DNA binding domain comprised of a helix-turn-helix motif (Korner *et al*., 2003, Soberon-Chavez *et al*., 2017). These transcriptional regulators have been shown to control several processes central to the lifestyle of their respective bacteria, such as carbon or nitrogen source utilization, aerobic/anaerobic transition, developmental processes, and pathogenicity (Lazazzera *et al*., 1993, Spiro & Guest, 1990, Derouaux *et al*., 2004, Kanack *et al*., 2006). Despite the presence of a common cNMP domain, individual groups within the family are regulated by diverse signals and transcriptional activity is controlled by different mechanisms. For example, the canonical *E. coli* Crp protein controls catabolite repression. Crp exists as an inactive dimer, which upon binding of cAMP results in allosteric reorientation of the DNA binding region allowing efficient binding to target DNA sequences (Saha *et al*., 2015), where it activates or represses downstream genes by making specific (Benoff *et al*., 2002). In contrast, for *E. coli* Fnr which induces genes necessary for anaerobic growth, activation is controlled by monomer to dimer transition (Lazazzera *et al*., 1993). Fnr senses oxygen via an associated Fe-S cofactor coordinated by four cysteine residues. In the absence of oxygen (activating conditions), the Fnr dimer is stabilized by a 4Fe-4S cluster and can bind to target sequences to activate or repress gene expression (Kiley & Beinert, 1998). In the presence of oxygen, the cluster transitions ultimately to an 2Fe-2S cluster leading to destabilization of the dimer and loss of DNA binding. In yet another variation, the *Xanthomonas campestris* CLP protein, which controls several genes involved in pathogenesis of plants, is a dimer which intrinsically binds target DNA sequences in the absence of ligand, but binding of di-c-GMP causes it to shift to an inactive conformation to release DNA binding sequences (Chin *et al*., 2010).

MrpC is a Crp/Fnr family member necessary for the starvation-induced multicellular developmental program of *Myxococcus xanthus* (Sun & Shi, 2001). *M. xanthus* is a gram negative deltaproteobacterium commonly found in the soil (Munoz-Dorado *et al*., 2016). In vegetative (non-developing conditions), *M. xanthus* is a cooperative predator. Swarms of *M. xanthus* cells glide in search of prey microorganisms or decaying organic material. Upon encountering prey, the swarm collectively releases antibiotics and degradative enzymes to paralyze and digest the prey. Upon nutrient poor conditions, the swarm enters a developmental program culminating in the formation of multicellular fruiting bodies filled with environmentally resistant spores. During this program, cells are first directed to move into haystack-shaped mounds (aggregation centers) of approximately 100,000 cells. Exclusively within these mounds, cells are induced to differentiate into spores, forming mature fruiting bodies. Production of spores inside fruiting bodies is not the only cell fate and accounts for only ∼15% of the starting population (as determined for the *M. xanthus* strain DZ2 induced to develop under submerged culture)(Lee *et al*., 2012). The majority of the cells (∼80%) undergo cell lysis, likely via programmed cell death (Rosenbluh *et al*., 1989, Wireman & Dworkin, 1977, Lee *et al*., 2012). The remaining ∼5% of cells are found as peripheral rods that remain outside of the fruiting bodies in a persister-like state (O’Connor & Zusman, 1991). Spore-filled fruiting bodies are quiescent and resistant to environmental insults, such as desiccation and UV, but upon return of nutrients, spores can germinate *en mass* to produce a productive feeding swarm (Munoz-Dorado *et al*., 2016).

MrpC plays a central role in the genetic regulatory network controlling the developmental program (Kroos, 2007), and an Δ*mrpC* mutant is incapable of aggregation and fails to launch the core sporulation program (Sun & Shi, 2001, McLaughlin *et al*., 2018). The *mrpC* gene is upregulated early after starvation and is dependent upon the MrpAB two component signal transduction system (Sun & Shi, 2001). MrpB is an enhancer binding protein, which when activated by phosphorylation of its associated receiver domain, induces transcription of *mrpC* from a putative sigma^54^-dependent promoter (Sun & Shi, 2001). Once produced, MrpC functions as a negative autoregulator, by competing with MrpB for overlapping binding sites in the *mrpC* promoter (McLaughlin *et al*., 2018). At least two other MrpC binding sites located upstream of the *mrpC* transcriptional start, also contribute to negative autoregulation [(McLaughlin *et al*., 2018) and unpublished data)].

MrpC is a global regulator, controlling at least 200 genes during the developmental program (Robinson *et al*., 2014). A key downstream target is the transcription factor, FruA (Ueki & Inouye, 2003, Ogawa *et al*., 1996). FruA is an orphan response regulator that is thought to induce aggregation and then sporulation in response to C-signaling, a cell-cell contact signal which increases in intensity as cells enter into aggregation centers (Saha *et al*., 2019, Ellehauge *et al*., 1998, Sogaard-Andersen & Kaiser, 1996). Activated FruA and MrpC act separately and in combination to regulate a number of genes necessary for differentiation of cells inside aggregation centers into spores (Son *et al*., 2011, Lee *et al*., 2011, Mittal & Kroos, 2009a, Mittal & Kroos, 2009b, Viswanathan *et al*., 2007).

It is unclear how MrpC binding to target promoters is controlled. No native ligand for MrpC is currently known, and the protein binds efficiently to target promoters *in vitro* (Ueki & Inouye, 2003, Nariya & Inouye, 2006, Mittal & Kroos, 2009a, McLaughlin *et al*., 2018) suggesting that MrpC is intrinsically able to binding target sequences in the absence of ligand. *In vivo*, MrpC is subject to several post-translational regulatory mechanisms. The EspAC signaling system functions early during development to activate an unknown protease to target MrpC to ensure only a gradual accumulation of MrpC that is presumably necessary for production of large, well-formed aggregation centers before the onset of sporulation (Schramm *et al*., 2012, Higgs *et al*., 2008, Cho & Zusman, 1999). In a second, apparently unrelated system, addition of nutrients to developing cells triggers unknown protease(s) to rapidly degrade MrpC allowing reversal out of the developmental program (prior to commitment to sporulation)(Rajagopalan & Kroos, 2014). Another post-translational regulation mechanism involves the serine/threonine protein kinase (STPK) cascade comprised of Pkn8 and Pkn14. *In vitro*, Pkn8 phosphorylates Pkn14, and Pkn14 phosphorylates MrpC on threonine residue(s) in its extreme amino terminus (Nariya & Inouye, 2005b, Inouye & Nariya, 2008, Nariya & Inouye, 2006). Phosphorylation was proposed to prevent LonD protease-dependent processing to remove the amino terminal 25 residues, producing ‘MrpC2’ (here termed MrpC_ΔN25_), a more active isoform. We have recently demonstrated, however, that MrpC’s amino terminal extension is essential for function *in vivo*, and that ‘MrpC2’ is likely an artifact of cell lysis (McLaughlin *et al*., 2018).

To reconcile the previous connection between phosphorylation of the MrpC amino-terminus with the recent demonstration that the amino terminus is essential for function, we set out to revisit Pkn14-dependent phosphorylation of MrpC. We demonstrate here that a specific cluster of threonines and serines (termed the TTSS motif) in the amino-terminal region is essential for MrpC activity *in vivo*. Alanine substitution of the complete MrpC TTSS motif prevents efficient development by interfering in MrpC’s negative autoregulation, proteolytic turnover, and transcriptional activation of FruA. Interestingly, while no single residue was necessary for activity, specific combinatorial substitutions within the TTSS motif produced highly variable developmental phenotypes revealing a previously unknown role for MrpC in developmental buffering. Mass spectrometry analysis of *in vitro* Pkn14-dependent phosphorylation of MrpC revealed that the MrpC TTSS motif is not directly phosphorylated by Pkn14 but is required for efficient phosphorylation of to several residues within the cNMP and DNA binding domains.

Reexamination of the role of Pkn8/14 suggests they are not active kinases during the developmental program of wild type *M. xanthus* under laboratory conditions, but likely fine tune MrpC activity in response to unknown environmental conditions. Thus, our data revises the model for Pkn8/14 control over MrpC activity and identifies an unusual TTSS motif that likely plays a role in stabilizing transitions between active and inactive MrpC states.

## RESULTS

### A TTSS motif within the N-terminal extension is necessary for MrpC activity

MrpC is a Crp/Fnr family transcriptional regulator with a 29 residue N-terminal extension that is essential for *in vivo* function (McLaughlin *et al*., 2018). No activating ligand is known for MrpC, but it was previously reported that MrpC activity is regulated by a STPK cascade which was proposed to phosphorylate MrpC on threonine residue(s) within the first 25 residues (Nariya & Inouye, 2005b, Nariya & Inouye, 2006). Sequence analysis of the amino-terminal region of MrpC shows that there are only two threonine residues (at positions 21 and 22) which are directly followed by two consecutive serine residues (Fig. 1A). As serine and threonine residues can both be phosphorylated by Ser/Thr kinases, we considered the TTSS residues a putative phosphorylation motif. To examine whether this motif was important for activity, we generated a strain in which the TTSS residues were entirely substituted with alanines in the endogenous *mrpC* locus (*mrpC_AAAA_*) and analyzed the resulting developmental phenotype under submerged culture conditions compared to the wild type and Δ*mrpC* strains. As expected, the wild type strain produced obvious aggregates between 34 and 48 hours which darkened by 72 hours of development, while the Δ*mrpC* strain failed to aggregate at all (Fig. 1B). The *mrpC_AAAA_* strain produced abnormally shallow and elongated aggregation streams that failed to progress to aggregation centers. Analysis of heat and sonication resistant spores at 72 hours of development indicated the wild type produced 2.6 ± 0.5 x 10^7^ heat and sonication resistant spores per well, while the Δ*mrpC* and *mrpC_AAAA_*mutants produced ≤ 0.01 % and 11 ± 10 % of wild type spore levels, respectively (Fig. 1C).

**Figure 1.**
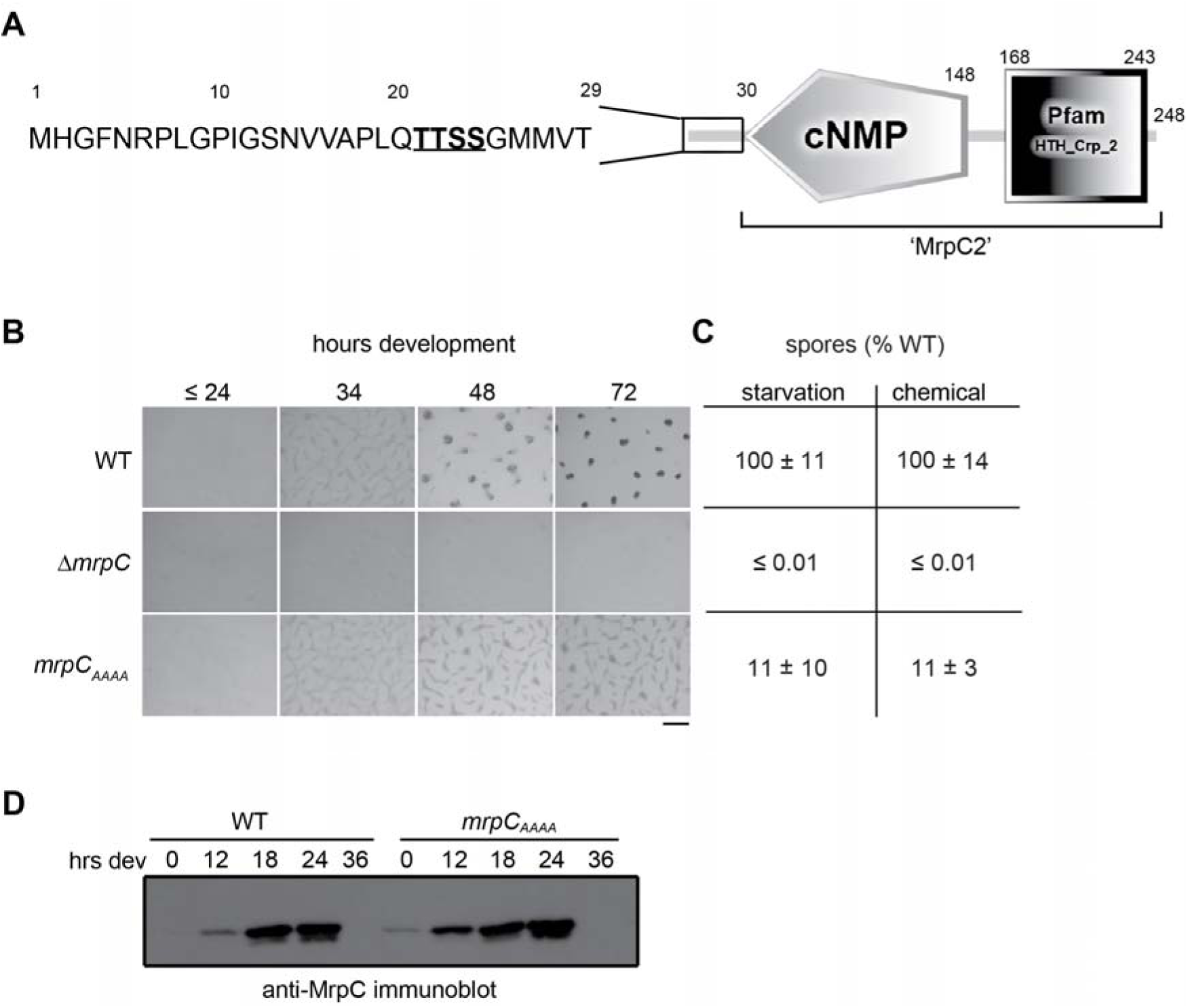
A TTSS motif within the N-terminus of MrpC is necessary for function. A. Domain architecture of MrpC depicted by SMART analysis (Letunic & Bork, 2018) showing the predicted cyclic nucleotide monophosphate binding (cNMP) and DNA binding domains (Pfam HTH_Crp_2). The sequence of the N-terminal 29 residues is shown with the TTSS motif underlined and in bold. Residue numbers are depicted above. “MrpC2” (aka MrpC_N25_) is a truncated MrpC that lacks the N-terminal 25 residues. B. Developmental phenotypes of wildtype (wt: DZ2), • *mrpC* (PH1025), and *mrpC_AAAA_* (PH1139) strains induced to develop under submerged culture and recorded at the indicated hours post-starvation. Bar: 0.5 mm C. Percent of wild type heat and sonication resistant spores harvested from cells developed for 72 hours under submerged culture (starvation) or after 24 hours of chemical induction with 0.5 M glycerol (chemical). Values are the average and associated standard deviations from three independent biological replicates. D. Anti-MrpC immunoblot analysis of protein lysates harvested from wild type (WT: DZ2) or *mrpC_AAAA_*(PH1139) cells developing under submerged culture for the indicated hours.

Since launch of the sporulation program is coupled to completion of aggregation, we could not distinguish whether defective sporulation by the *mrpC_AAAA_* mutant was because it failed to complete aggregation or whether MrpC_AAAA_ specifically interfered in induction of the sporulation program. To distinguish between these two possibilities, we examined the sporulation efficiency of wild type, Δ*mrpC*, and *mrpC_AAAA_* strains upon chemical induction of sporulation which bypasses the requirement for aggregation (Dworkin & Gibson, 1964). For this assay, vegetative broth cultures of wild type, Δ*mrpC*, or *mrpC_AAAA_* cells were treated with 0.5 M glycerol for 24 hours and heat and sonication resistant spores were counted. We observed that the Δ*mrpC* and *mrpC_AAAA_* strains produced ≤ 0.01 and 11 ± 3 % of wildtype spore levels, respectively (Fig. 1C), suggesting that MrpC_AAAA_ was strongly reduced in triggering spore differentiation.

To determine whether the *mrpC_AAAA_* defective developmental and sporulation phenotypes were simply explained by reduction in MrpC stability, we compared the levels of MrpC produced from the wild type, and *mrpC_AAAA_* strains at 0, 12, 18, 24, and 36 hours of development by anti-MrpC immunoblot. As expected, wild type MrpC protein began to accumulate at 12 hours but was absent by 36 hours (Fig. 1D). The MrpC_AAAA_ protein had ∼ 2-fold increased accumulation relative to the parent strain at 12 hours but was also absent by 36 hours development (Fig 1D). Thus, the development defect observed in by the *mrpC_AAAA_*mutant was not due to failure to accumulate MrpC_AAAA_, strongly suggesting that an intact TTSS motif is necessary for MrpC function.

### Consecutive intact residues within the MrpC TTSS motif are necessary for robust development

To determine whether substitution of any specific residue within the TTSS motif was sufficient to observe the *mrpC*_AAAA_ phenotype, strains bearing individual alanine substitutions of each TTSS residue were generated in the endogenous *mrpC* locus. These strains, producing MrpC_T21A,_ MrpC_T22A_, MrpC_S23A_ or MrpC_S24A_ are hereafter designated with the TTSS substitution (i.e. *mrpC*_ATSS_, *mrpC*_TASS_, *mrpC*_TTAS_ or *mrpC*_TTSA_, respectively). These mutants produced developmental phenotypes and sporulation efficiencies similar to the wild type strain (Fig. S1), suggesting no one particular individual residue in the TTSS motif was necessary for activity. Furthermore, substitution of both threonines or both serines of the TTSS motif in the endogenous *mrpC* gene (strains *mrpC*_AASS_ or *mrpC*_TTAA_), respectively) also produced a wild type phenotype suggesting the neither of the threonine nor the serine residues were functionally redundant (Fig. S1).

We next considered whether any intact single residue within the TTSS motif was sufficient for function by constructing two different mutants in which only one residue was available for phosphorylation: *mrpC*_TAAA_ or *mrpC*_AAAS_. As these mutants proved difficult to generate in the endogenous *mrpC* locus, we instead expressed the *mrpC*_TTSS_ (wt), *mrpC*_AAAA_, *mrpC*_TAAA_, or *mrpC*_AAAS_ clones from their endogenous promoter inserted at the Mx8 phage attachment (*attB*) site in the Δ*mrpC* background (Δ*mrpC attB*::P_mrpC_-*mrpC*_TTSS_, etc.; termed *att*::*mrpC*_TTSS,_ etc. for short). The *att*::*mrpC*_TTSS_ (wt) clone complemented Δ*mrpC*, although aggregation onset was observed on average four hours earlier than the DZ2 parent (data not shown). The *att*::*mrpC*_AAAA_ mutant phenocopied *mrpC*_AAAA_ generated at the endogenous locus (Fig. 1 and Fig. 2). Strikingly however, independent clones and replicates of the same clones of the *att*::*mrpC*_TAAA_ or *att*::*mrpC*_AAAS_ strains did not display stable developmental phenotypes. To quantify the extent of phenotypic variation between replicates and between clones, we used a high-throughput, high resolution development imaging technique to generate movies of strains undergoing development (Glaser & Higgs, 2019). Stages of development were recorded for each movie and displayed as heat maps versus the indicated ranges of hours post-starvation (Fig. 2). In this assay, wild type *mrpC* clones displayed robust developmental phenotypes, with onset of aggregation observed between 24-29.5 hours post starvation, followed by initial aggregates (30-35.5 h), aggregates after consolidation/dissolution (36-41.5 h), mature immobile aggregates (42-47.5 h) and darkened fruiting bodies (≥ 48 h). The *att*::*mrpC*_AAAA_ clones consistently failed to develop properly (Fig. 2). However, for the *att*::*mrpC*_TAAA_ or *att*::*mrpC*_AAAS_ strains, significantly different developmental patterns were observed both among clones and between replicates of the same clone (Fig. 2). Developmental patterns observed ranged from early or delayed development to failure to develop (Fig. 2). We observed the same variability in phenotypes when the *mrpC* variants were instead integrated at a different secondary site [termed 1.38kb (Garcia-Moreno *et al*., 2010)] in the Δ*mrpC* background, indicating that the variability was not due to placement at the *attB* site (Fig. S2). These results suggested that induction of development had become stochastic, perhaps because the equilibrium between active and inactive MrpC was perturbed.

**Figure 2.**
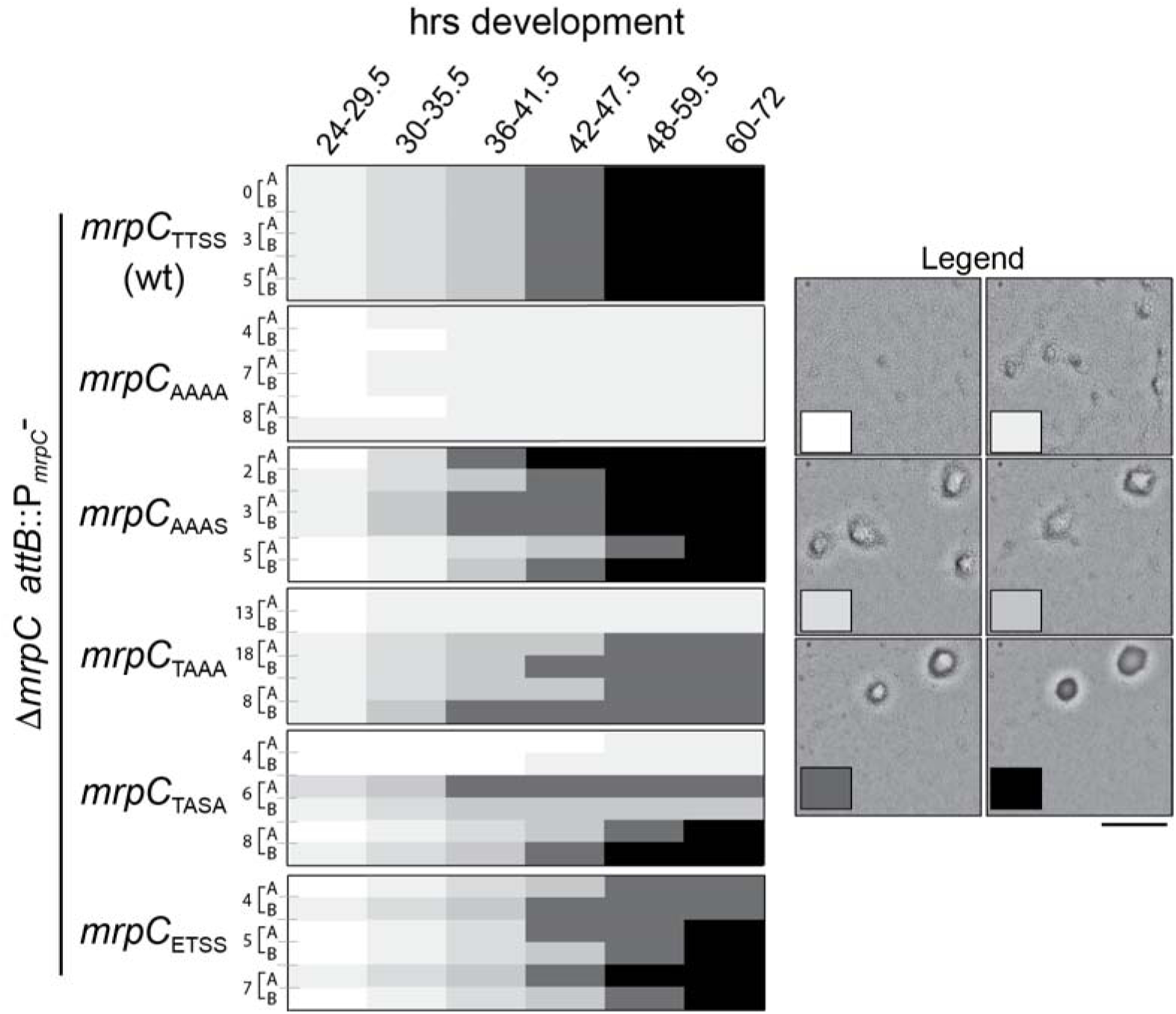
Lack of consecutive intact MrpC TTSS motif residues leads to loss of developmental robustness. Inter- and intra-clone variability in the developmental phenotypes of *mrpC*_TTSS_ (wild type)(PH1118), *mrpC_AAAA_* (PH1530), *mrpC_AAAS_*(PH1532), *mrpC_TAAA_* (PH1531)*, mrpC_TASA_* (PH1547), and *mrpC*_ETSS_ (PH1533) genes integrated at the *attB* site in the Δ*mrpC* background. The heat maps shown indicate the variability in the timing and extent of completion of fruiting body formation during development under submerged culture conditions in 96 well plates in the indicated ranges of developmental times. Replicates are indicated by letters (A,B), and independent clones are represented by clone number. The shade of box corresponds to the extent of development as indicated by representative pictures in the legend (see text for details). Bar: 250 µm.

Assuming the TTSS motif was indeed necessary for phosphorylation, we postulated that since the *mrpC*_AASS_ and *mrpC*_TTAA_ mutants displayed stable wild type phenotypes (Fig. S1), perhaps any two intact TTSS residues were necessary for MrpC function. To test this hypothesis, we generated *att*::*mrpC*_TASA_ clones and again observed that this mutant produced variable developmental phenotypes both between clones and replicates of the same clone (Fig. 2). Together, these results suggested that at least two consecutively intact TTSS residues were necessary for MrpC function. However, from these data it was not fully clear whether the TTSS motif was directly phosphorylated or whether it served as an important polar motif necessary for stable conformational switching or protein-protein interactions.

Finally, as an alternate genetic approach, we attempted to generate phosphomimetic mutations in which substitution with glutamic acid may mimic a phosphorylated amino acid (Dissmeyer & Schnittger, 2011). For these analyses, we generated *att*::*mrpC*_ETSS_, *att*::*mrpC*_EAAA_, or *att*::*mrpC*_EEAA_ clones to examine whether “forcing” at least one phosphorylation but leaving the remaining motif intact, or generating “constitutive single only” or “constitutive double” phosphorylation states, could reveal a phospho-code relating to function. Analysis of the developmental phenotype of these mutants indicated the ETSS substitution produced slightly variable, delayed to wild type aggregation phenotypes with fruiting bodies that sometimes failed to darken (Fig. 2), which usually indicates impaired sporulation. Both the *att*::*mrpC*_EAAA_ and *att*::*mrpC*_EEAA_ mutants were essentially inactive and phenocopied the *att*::*mrpC*_AAAA_ phenotype (data not shown). These results suggested either that glutamic acid substitution did not act as a phosphomimetic or that the TTSS motif is not a phosphorylation target.

### Pkn14 phosphorylates MrpC in vitro on several residues in the cNMP and DNA-binding domains

To determine whether we could recapitulate the observation that Pkn14 phosphorylates MrpC *in vitro* (Nariya & Inouye, 2005b), we overexpressed and purified Strep affinity tagged-Pkn14 and a kinase-dead version of the protein in which the conserved lysine at position 48 which is predicted to be necessary for ATP binding (Hanks, 2003) was substituted with asparagine (Strep-Pkn14_K48N_). Strep-Pkn14 or Strep-Pkn14_K48N_ was incubated for 30 minutes in the presence of ATP, resolved by SDS-PAGE, and the resulting gel was incubated in phosphoprotein stain. Autophosphorylated Strep-Pkn14 could be detected readily, while the signal on Pkn14_K48N_ was significantly reduced (Fig. 3A); the remaining signal was likely due to non-specific fluorescence because the Pkn14_K48N_ protein was completely inactive when incubated in the presence of [γ-^32^P] ATP (data not shown). We next repeated these assays in the presence of purified hexa-histidine affinity tagged (His_6_) -MrpC, -MrpC lacking the 25 amino terminal residues (His_6_-MrpC_ΔN25_), His_6_-MrpC_AAAA_, or the non-specific protein Trx-His_6_. Phosphorylation of the MrpC protein could be detected, which was reduced 1.7- and 4.0-fold on His_6_- MrpC_ΔN25_ or His_6_-MrpC_AAAA_, respectively. Some signal was observed on the non-specific protein Trx-His_6_ (Fig. 3A). Thus, we recapitulated the result that Pkn14 appears to phosphorylate MrpC and show that phosphorylation is reduced (but not absent) if the TTSS motif is removed.

**Figure 3.**
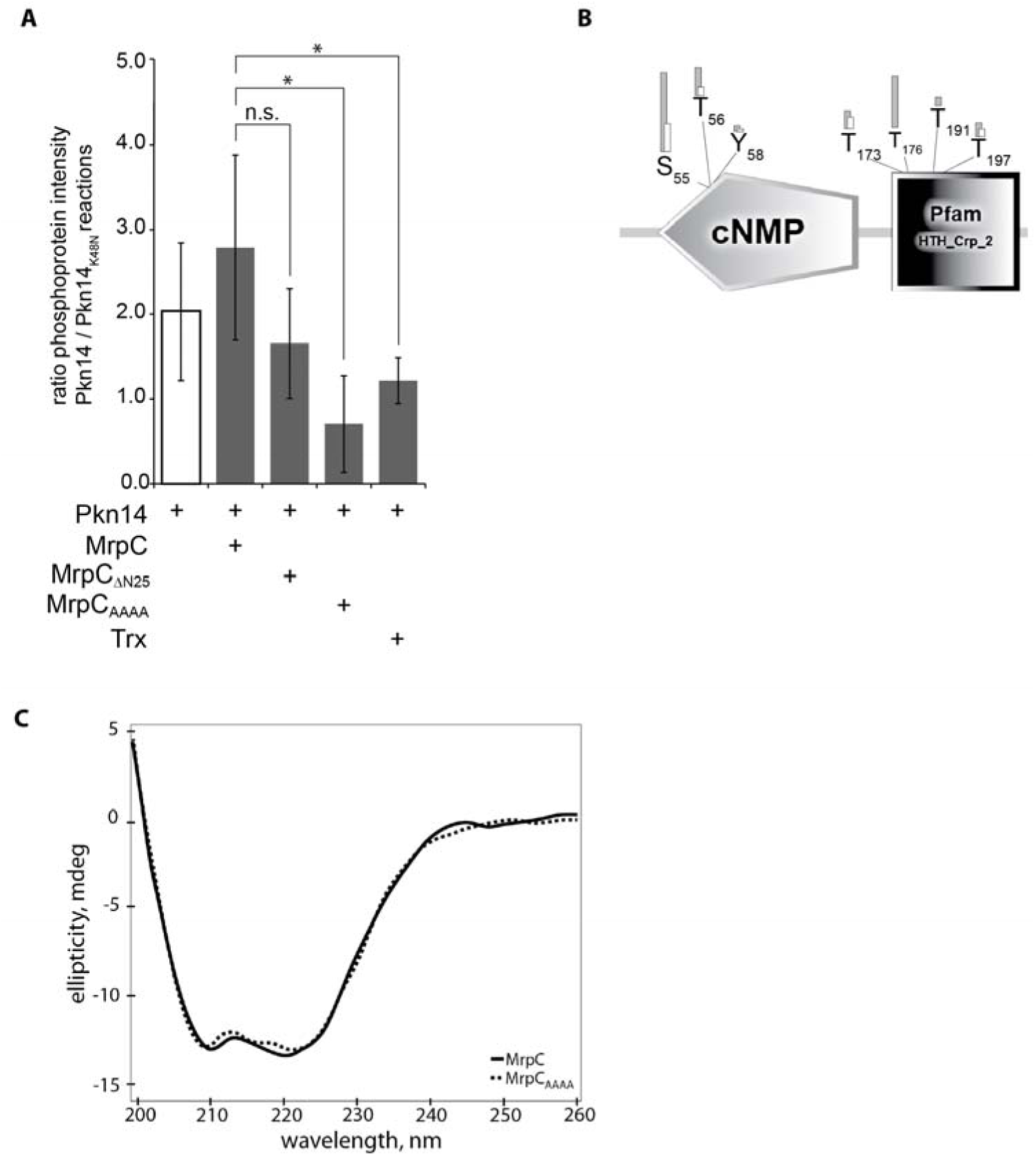
Pkn14 does not phosphorylate MrpC on the TTSS motif *in vitro*. A. Specific phosphoprotein stain intensities from Strep-Pkn14 autophosphorylation (white bar) or from Pkn14-dependent phosphorylation of His_6_-MrpC, His_6_-MrpC_ΔN25_, His_6_-MrpC_AAAA_, or the non-specific protein Trx-His_6_, as indicated (grey bars). For each reaction, Strep-Pkn14 or –Pkn14_K48N_ were incubated in kinase buffer and indicated substrate. Reactions were resolved by protein gel electrophoresis, and phosphorylated proteins were detected by phosphoprotein stain, and then by total protein stain. Phosphoprotein signal intensities were first normalized to total protein and then normalized to phosphoprotein signal intensities from the kinase dead Pkn14_K48N_ reactions. Data points are the averages and associated standard deviations from three independent replicates. *, *p*-value ≤ 0.05 as determined by Student’s t-test; n.s., not significant. B. Pkn14-dependent phosphorylated residues on MrpC detected by mass spectrometry. *In vitro* kinase reactions containing Pkn14 or Pkn14_K48N_ and MrpC or MrpC_AAAA_ were trypsin digested and TiO_2_-enriched phosphopeptides were subject to MS/MS analysis. Peptides are shown which meet stringent cutoff criteria (minimum 5 peptides detected, 0.1% FDR). Phosphorylation counts on MrpC (grey bars) or MrpC_AAAA_ (white bars) residues are shown. No phosphorylation counts were detected in the respective Pkn14_K48N_ reactions. The MrpC domain architecture from SMART analysis is depicted (Letunic & Bork, 2018). C. Circular dichroism analysis of His-MrpC (blue trace) and His-MrpC_AAAA_ (red trace). Measurements were performed under ionic conditions similar to those used in our *in vitro* assays.

We next extended these analyses to examine which residues in His_6_-MrpC or His_6_-MrpC_AAAA_ might be phosphorylated by Strep-Pkn14 *in vitro* using mass spectrometry. For this approach, Strep-Pkn14 or Strep-Pkn14_K48N_ were incubated with His_6_-MrpC or His_6_-MrpC_AAAA_ using the *in vitro* phosphorylation reaction conditions. Reactions were quenched, trypsin digested, and phosphopeptides were captured on titanium dioxide columns, separated by liquid chromatography and subjected to mass spectrometry. In the Pkn14/MrpC reaction, we captured 72 phosphopeptides. With strict selection criteria (minimum 5 peptides detected, 0.1% false discovery rate), Ser/Thr phosphoresidues detected corresponded to MrpC S_55_, T_56_ (within the cNMP-binding domain), and T_173_, T_176,_ T_191_, and T_197_ (within the DNA binding domain)(Fig. 3B). These phosphorylated residues were observed in two independent replicates, although the number of phosphopeptides captured varied slightly. In the Pkn14/MrpC_AAAA_ reaction, we observed only 20 phosphopeptides, but they corresponded to the same sites as the wild type with the exception of T_176_ and T_191_. As expected, no phosphopeptides could be detected in the Pkn14_K48N_/MrpC or Pkn14_K48N_/MrpC_AAAA_ reactions, indicating that all phosphopeptides observed in the wild type Pkn14 reactions were a result of Pkn14 specifically. Thus, the TTSS motif was not in fact phosphorylated by Pkn14 *in vitro* suggesting the reduction in phosphorylation on the MrpC_AAAA_ was likely a result of inefficient recognition of MrpC as a substrate by Pkn14.

To rule out that the conformation of His_6_-MrpC_AAAA_ was drastically perturbed, we compared the circular dichroism (CD) spectra of His_6_-MrpC and His_6_-MrpC_AAAA_ under ionic conditions similar to those used in our *in vitro* assays (Fig. 3C). The secondary structures of His_6_-MrpC_AAAA_ and His_6_-MrpC were identical, with characteristic absorbance peaks at λ = 208 and 220 nm. Thus, substitution of the TTSS motif with alanines did not produce drastic changes in the secondary structure of MrpC. Together, these results suggested that the TTSS motif may instead be a MrpC recognition motif.

### The Pkn8/Pkn14 kinase cascade does not inactivate MrpC in the wild type strain under laboratory conditions

Regardless of whether the TTSS motif is directly phosphorylated by Pkn14 or is merely a recognition motif that facilitates Pkn14-dependent phosphorylation of MrpC at different sites, the observation that the *mrpC*_AAAA_ mutant does not develop suggested that phosphorylation of MrpC would serve as an activation signal, rather than the inactivation signal originally proposed based on the early developmental phenotypes previously observed for both *pkn14* and *pkn8* mutant strains in the non-wildtype *M. xanthus* background strain, DZF1 (Nariya & Inouye, 2005b). As the DZF1 (aka DK101) strain is defective in a social motility system (Wall *et al*., 1999) which can often perturb developmental phenotypes (Lee *et al*., 2012, Boynton *et al*., 2013), we next sought to reexamine the developmental phenotypes of strains lacking the Pkn8/Pkn14 kinase cascade in our wild type DZ2 *M. xanthus* background. We therefore generated in-frame deletions of *pkn14* and *pkn8* and constructed point mutations predicted to render each protein kinase-dead in the endogenous locus of each gene (strains Δ*pkn14*, Δ*pkn8*, *pkn14*_K48N_, *pkn8*_K116N_, respectively). When these strains and the wild type were induced to develop under submerged culture conditions, the wild type strain produced visible aggregation centers between 28- and 35-hours post-starvation, and 2.7 ± 0.7 x 10^7^ heat and sonication resistant spores at 120 hours (Fig. 4A). In contrast to previously published results, the Δ*pkn14* strain showed delayed aggregation: visible aggregation centers were detected between 35 and 48 hours, approximately 6 hours later than the wild type. By 120 hours of development, the Δ*pkn14* mutant sporulated at wild type efficiencies (102 ± 15 % of wild type)(Fig. 4A and B). When we generated our Δ*pkn14* in the DZF1 strain background and developed the strains on nutrient limited CF agar (DZF1 strains do not develop under submerged culture), we observed the same early developmental phenotype originally observed (Nariya & Inouye, 2005b), while the DZ2 Δ*pkn14* strain again exhibited a delayed developmental phenotype (Fig. S3). Thus, we concluded that the difference in developmental phenotypes was not the result of different Δ*pkn14* constructs or developmental conditions assayed, but rather due to strain background. We and others have previously observed that the DZF1 strain background yields different mutant phenotypes compared to the wild type strains DZ2 and DK1622 (Lee *et al*., 2012, Boynton *et al*., 2013). These results in the wild type background were consistent with a model in which Pkn14-dpendent phosphorylation could activate MrpC.

**Figure 4.**
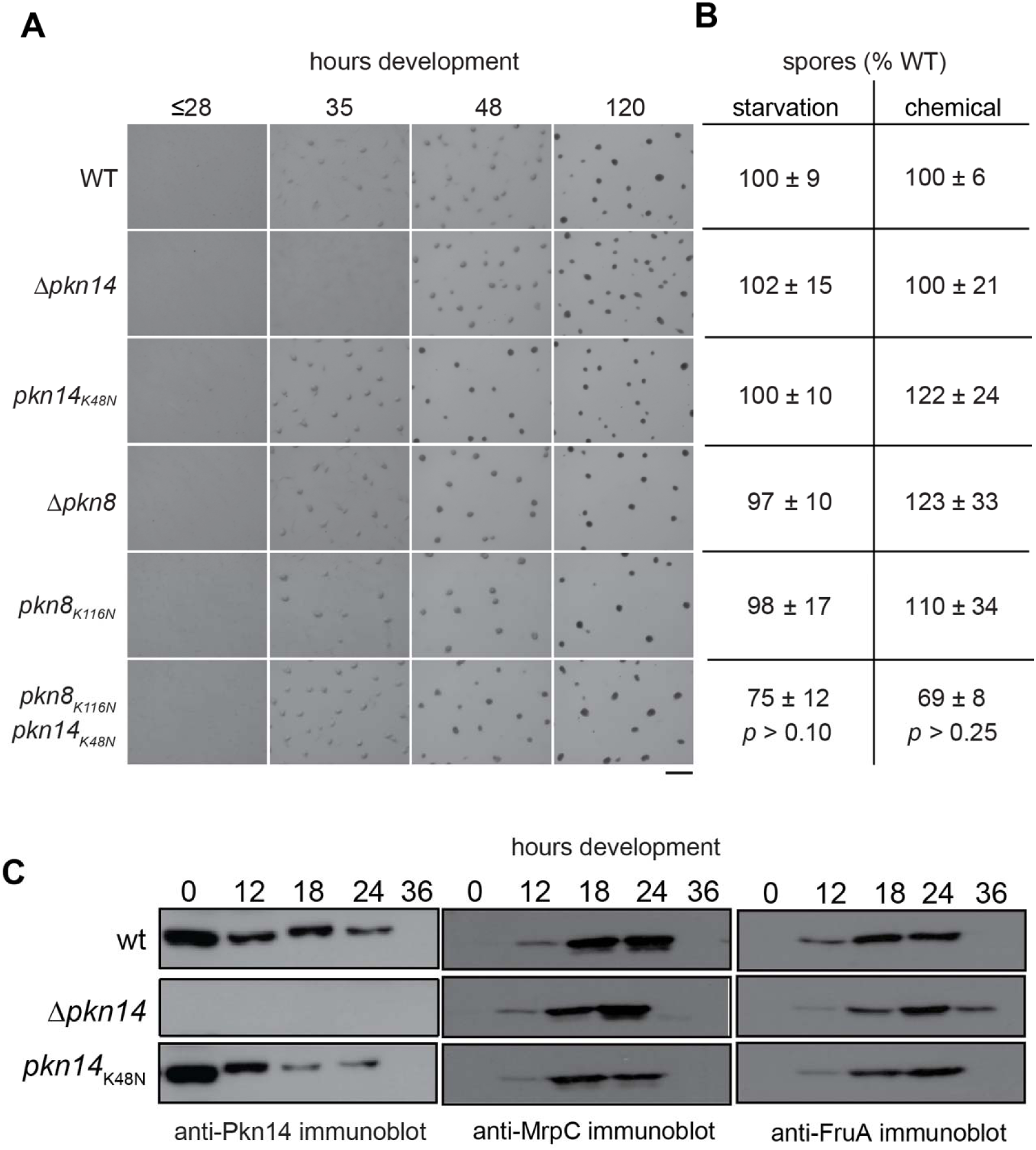
Pkn8/Pkn14 do not repress development in the *M. xanthus* DZ2 strain background. A. Developmental phenotypes of wild type DZ2, Δ*pkn14* (PH1132), *pkn14_K48N_* (PH1133), Δ*pkn8* (PH1347), *pkn8_K116N_* (PH1548) and double point mutant *pkn14_K48N_ pkn8_K116N_* (PH1549) strains were induced to develop by submerged culture conditions and images were recorded at the indicated hours of development. Bar: 0.5mm. B. Sporulation efficiencies from starvation-induced development (left) and chemical induced development (right). The number of heat and sonication resistant spores harvested after 120 hrs development (left) or 24 hours of induction (right) and reported as a percent of wildtype, respectively. Values are the average and associated standard deviations from three independent biological replicates. *p*-value was determined by chi-square analysis. C. Pkn14, MrpC, and FruA accumulation profiles in wild type (DZ2), Δ*pkn14* (PH1132), *pkn14_K48N_* (PH1133) or cells. 10 µg lysates generated from cells induced to develop under submerged culture for the indicated hours post-starvation were subject to anti-Pkn14, -MrpC, or -FruA immunoblot, as indicated.

Surprisingly, however, analysis of the *pkn14*_K48N_ mutant indicated this strain produced a developmental phenotype indistinguishable from the wild type with respect to fruiting body production and sporulation efficiency (100 ± 10 % of wild type). It is unlikely that the K48N substitution did not inactivate Pkn14 kinase activity, because it rendered Pkn14 incapable of autophosphorylation *in vitro* (Fig. 3A and data not shown). To confirm the mutant protein accumulated properly during development, we generated antibodies against the Pkn14 protein and performed anti-Pkn14 immunoblot analysis on protein lysates harvested from wild type, Δ*pkn14* and *pkn14_K48N_* mutants at 0, 12, 18, 24, and 36 hours of development. A band migrating at ∼48 kDa could be detected in the wild type, but not Δ*pkn14* lysates, consistent with the Pkn14 predicted molecular mass of 45.4 kDa (Fig. 4C). Pkn14 was detected at highest levels in vegetative cells which decreased 7.2-fold by 12 hours, remained constant at 18 and 24 hours, but was absent by 36 hours. A similar pattern could be detected for Pkn14_K48N_ (Fig. 4C). Thus, Pkn14 was expressed until at least the onset of aggregation, and disruption of auto-phosphorylation did not significantly alter its accumulation pattern. To next examine whether the *pkn14* mutants exhibited perturbed MrpC or FruA accumulation, we probed the same samples with anti-MrpC or anti-FruA immunosera. We observed similar MrpC and FruA accumulation in both the mutant *pkn14* strains compared to the wild type, with the exception that FruA levels were very slightly delayed in the Δ*pkn14* mutant, likely as a result of the delayed development observed in this strain (Fig. 4C).

Analysis of the of the Δ*pkn8* and *pkn8_K116N_* mutants revealed that they produced developmental phenotypes indistinguishable from the wild type with respect to fruiting body formation (Fig. 4A) and sporulation efficiency (97 ± 10 % and 98 ± 17 % of wild type spores, respectively) (Fig. 4B); this phenotype was also in contrast to the early developmental phenotype observed in the DZF1 background (Nariya & Inouye, 2005b). Finally, to examine whether the Pkn14 and Pkn8 kinase activity was redundant, we generated a double *pkn14_K48N_ pkn8_K116N_* mutant. This mutant also displayed a wild type developmental phenotype (Fig. 4A). The small reduction in sporulation efficiency observed in this mutant during starvation (75 ± 12 %)- and glycerol (69 ± 8 %)-induced sporulation was not considered statistically significantly different from wild type (Fig. 4B).

In summary, our genetic analyses suggested that under our laboratory developmental conditions, Pkn8 has no obvious role in development and that the presence of Pkn14, but not its kinase activity, is necessary to promote efficient developmental aggregation.

These data suggested that Pkn14-dependent phosphorylation of MrpC likely plays a role in controlling MrpC in response to perturbed conditions observed in the DZF1 background, which we are currently investigating. However, as the dramatic *mrpC*_AAAA_ developmental phenotype suggested that the TTSS motif was essential for MrpC function, and little is known about how the activity of MrpC is intrinsically controlled, we set out to examine which of the activities attributed to MrpC were perturbed by this mutant.

### Perturbation of the MrpC TTSS motif perturbs in vivo MrpC activities

As MrpC functions as a negative autoregulator (McLaughlin *et al*., 2018) and the MrpC_AAAA_ protein accumulated slightly earlier than wild type (Fig. 1D), we first addressed whether negative autoregulation was perturbed. The effect on autoregulation was analyzed with a reporter containing mCherry (mCh) under control of the *mrpC* promoter (P_mrpC_-mCh) (McLaughlin *et al*., 2018) which was integrated at the Mx8 phage *att* site in the wild type, *mrpC*_AAAA_, and Δ*mrpC* backgrounds. The developmental phenotype observed in these strains bearing the reporter was indistinguishable from the parent strains (data not shown). To examine reporter activity, each strain was harvested at 0, 12, 18, 24, 30, 36, and 48 hours of development under submerged culture, and mCherry fluorescence was recorded and normalized to total protein. As we previously reported (McLaughlin *et al*., 2018), mCherry signal in the wild type P_mrpC_-mCh background gradually increased over 48 hours of development, and consistent with negative autoregulation, reporter activity increased at least 3.1-fold over wild type in the Δ*mrpC* P_mrpC_-mCh strain (Fig. 5A). In contrast, reporter activity in the *mrpC*_AAAA_ P_mrpC_-mCh strain was only slightly higher with a maximum fold induction of 1.9-fold over wild type at 36 hours (Fig. 5A). These results suggested that MrpC_AAAA_ has a slight defect in autoregulation which likely explains the elevated levels of MrpC_AAAA_ observed in the initial immunoblot analysis (Fig. 1).

**Figure 5.**
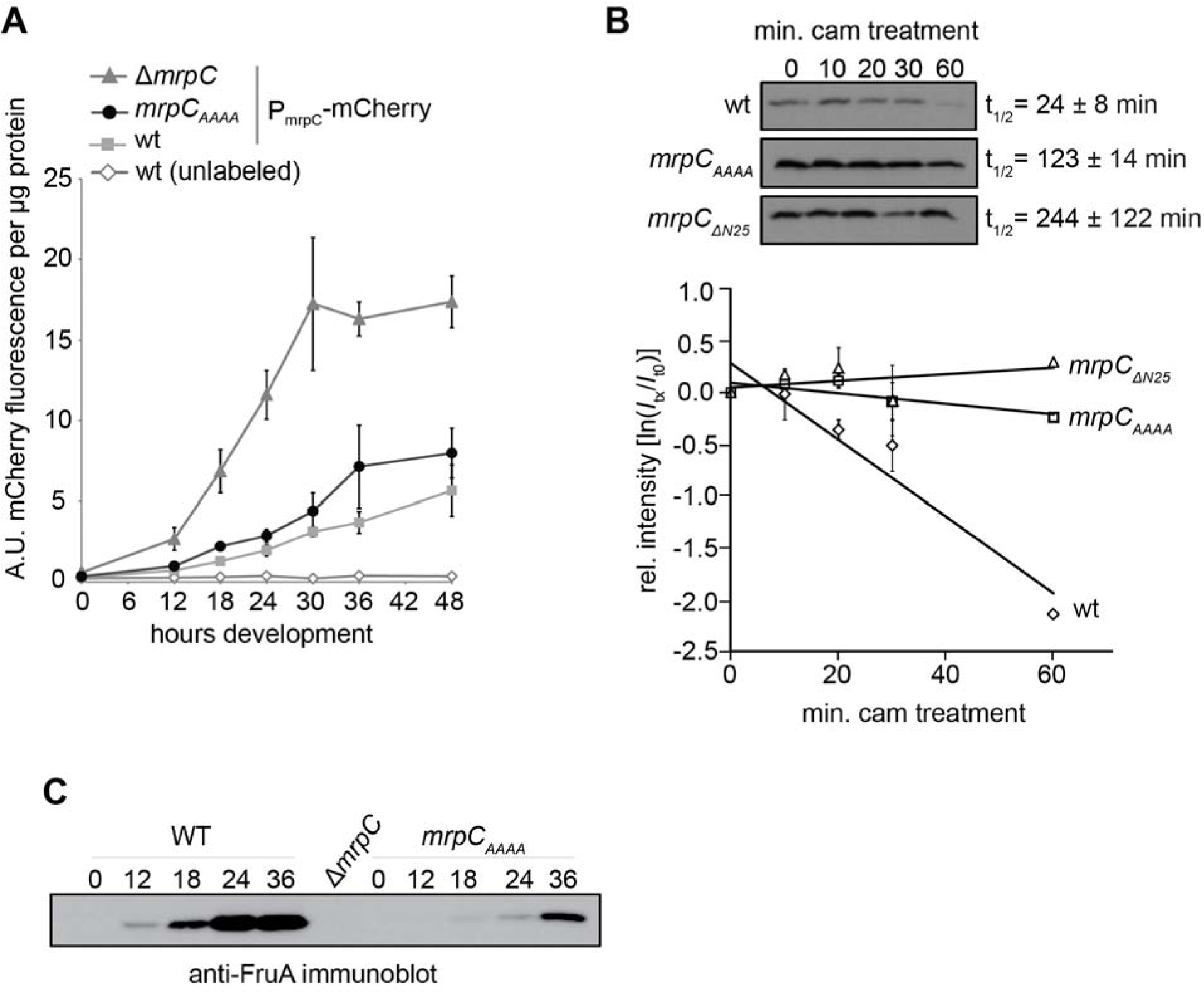
MrpC_AAAA_ is perturbed in *mrpC* negative autoregulation, MrpC turnover, and FruA induction. A. Expression of a P*_mrpC_-mCherry* reporter in wild type (PH1100), Δ*mrpC* (PH1104), and *mrpC_AAAA_* (PH1306) backgrounds. Cells were induced to develop under submerged culture conditions and harvested at the indicated times. mCherry fluorescence was recorded as arbitrary units (A. U.) and then normalized to total protein. Values are the average and associated standard deviations from three independent biological replicates. Wild type unlabeled (DZ2) cells lack the reporter. B. MrpC turnover assay. Wild type (wt; DZ2), *mrpC*_AAAA_ (PH1139), or *mrpC*_ΔN25_ (PH1108) strains were induced to develop under submerged culture conditions for 9 hours and treated with 34 ug ml^-1^ chloramphenicol (cam) for the indicated times. Equal cell amounts of each sample were subjected to anti-MrpC immunoblot (upper panels). MrpC half-lives were calculated from two independent biological replicates where the MrpC band intensity for each time point was normalized to the intensity at T=0 of the respective strain and, and the natural log of the normalized intensities was plotted versus min of chloramphenicol treatment (lower graph). The slope of the linear fit of the data was used to calculate the MrpC half-life (t_1/2_) in each strain for each replicate, and those values were averaged. Values plotted are the average and associated standard deviation for each time point from the two replicates. C. Anti-FruA immunoblot analysis of protein lysates harvested from Δ*mrpC attB*::P_mrpC_-*mrpC_TTSS_* (WT: PH1118) or DZ2 *mrpC_AAAA_*(PH1139) strains developing under submerged culture conditions for the indicated hours. The • *mrpC* lysate (PH1025) was prepared from cells at 24 hours of development.

During development, MrpC protein levels are also controlled by EspAC-dependent proteolytic turnover (Schramm *et al*., 2012). To examine whether the MrpC_ΔN25_ and MrpC_AAAA_ proteins were efficiently turned over, we examined the half-life of each protein in chloramphenicol shutoff assays. For these experiments, the wild type, *mrpC*_AAAA_, and *mrpC*_ΔN25_ strains were induced to develop under submerged culture conditions for nine hours, treated with chloramphenicol, and protein lysates harvested from cells after 0, 10, 20, 30, and 60 min were subject to anti-MrpC immunoblot. Consistent with previous observations (Schramm *et al*., 2012), we calculated an MrpC half-life of 24 ± 8 min. However, MrpC_AAAA_ and MrpC_ΔN25_ were not efficiently turned over (t_1/2_ = 123 ± 14 and 244 ± 122 min, respectively)(Fig 5B). These results suggest that MrpC_ΔN25_ and MrpC_AAAA_ are not efficiently targeted for regulated proteolysis which likely also contributes to the elevated levels observed *in vivo* for MrpC_AAAA_ (Fig.1D) or MrpC_ΔN25_ (McLaughlin *et al*., 2018).

MrpC is also a transcriptional activator and an important target is *fruA* (Ueki & Inouye, 2003). FruA is essential for induction of aggregation and sporulation (Ellehauge *et al*., 1998, Ogawa *et al*., 1996). We next examined the accumulation of FruA in the *att*::*mrpC* and *mrpC*_AAAA_ strains by anti-FruA immunoblot analysis of lysates prepared from cells induced to develop under submerged culture conditions for 0, 12, 18, 24 and 36 hours. In *att*::*mrpC* cells, FruA protein could be detected by 12 hours of development and continued to accumulate to at least 36 hours (Fig. 5C). No FruA could be detected in the Δ*mrpC* strain (Fig. 5C and data not shown). In contrast, production of FruA was severely delayed and reduced in the *mrpC*_AAAA_ lysates (Fig. 5C). Together, these results suggested MrpC_AAAA_ partially failed in repressing its own expression and was strongly impaired in activating *fruA* expression. The failure to efficiently induce FruA likely explained the failure to induce proper development observed in the *mrpC*_AAAA_ mutant (Fig. 1B).

### Perturbation of the MrpC TTSS motif reduces affinity for binding sites within the mrpC and fruA promoters

To examine whether the MrpC_AAAA_ protein was defective in binding to target promoters *in vitro*, we used electrophoretic mobility shift assays to determine the relative affinity of purified His_6_-MrpC or His_6_-MrpC_AAAA_ for fluorescently labeled probes containing an MrpC binding site (corresponding to -237 to -266 bp upstream from the *mrpC* start; aka BS 5)(McLaughlin *et al*., 2018). The bound and unbound fluorescent DNA probes were resolved by gel electrophoresis and probes were detected by fluorescence imager. As seen previously (McLaughlin *et al*., 2018), 0.5-2 μM His_6_-MrpC caused a progressively increasing shift in the *mrpC* binding site probe. No shift was observed when excess unlabeled probe was added indicating binding was specific (Fig 6A). In contrast, nearly 2 μM His_6_-MrpC_AAAA_ was required to detect a shift in the mobility of the probe, suggesting an approximately 4-fold reduction in binding affinity (Fig. 6A).

**Figure 6.**
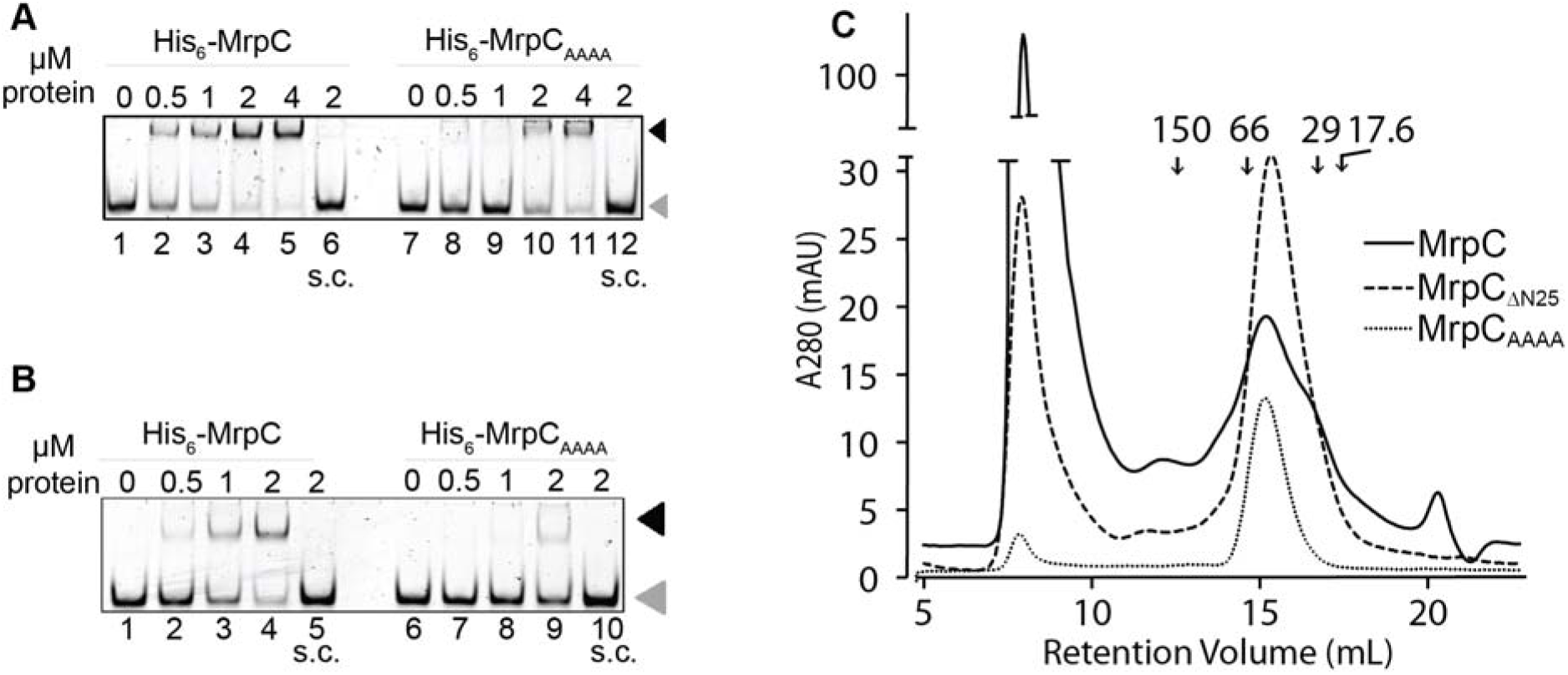
MrpC_AAAA_ does not efficiently bind to *mrpC* or *fruA* promoter binding sites. A. Electrophoretic mobility shift assays (EMSAs) of His_6_-MrpC or His_6_-MrpC_AAAA_ using *mrpC* promoter binding site 5. Increasing concentration (0-4 micromolar, as labeled) of His_6_-MrpC (lanes 1-6) or His_6_-MrpC_AAAA_ (lanes 7-12) was incubated with 50 nM of fluorescently labeled probe. B. EMSA as in A. using *fruA1* promoter binding site. Increasing concentration (0-2 micromolar, as labeled) of His_6_-MrpC (lanes 1-5) or His_6_- MrpC_AAAA_ (lanes 6-10) was incubated with 50 nM of fluorescently labeled probe. Lanes 6 and 12 (A), or 5 and 10 (B) additionally contain 2.7 µM of the respective unlabeled probe (s.c., specific chase). Black arrowhead, protein + DNA complex; gray arrowhead, unbound DNA probe. C. MrpC_AAAA_ and MrpC_ΔN25_ retain dimerization capability in solution. Analytical size-exclusion chromatography on a Superdex^TM^ 200 10/300 GL column using 4 nmol (8 µM x 500 µl) His_6_-MrpC (red), His_6_- MrpC_ΔN25_ (blue) or His_6_- MrpC_AAAA_ (black). Retention volumes of alcohol dehydrogenase (150 kDa), albumin (66 kDa), carbonic anhydrase (29 kDa), and myoglobin (17.6 kDa) are depicted. Peak MrpC, MrpC_ΔN25_, or MrpC_AAAA_ fractions corresponded to peaks at retention volumes of 15.3, 15.5, and 15.2 mL, respectively, consistent with MrpC dimers (∼50∼60 kDa).

To examine MrpC-binding to the *fruA* promoter, we first identified two putative MrpC binding sites situated at -366 to -395 (*fruA1*) and -341 to -370 bp (*fruA2*) from the *fruA* transcriptional start site, based on MrpC foot printing observed previously (Ueki & Inouye, 2003). When His_6_-MrpC was incubated with the *fruA1* probe, we observed a shift in probe migration with at least 0.5 μM His_6_-MrpC, whereas in the presence of His_6_- MrpC_AAAA_, an equivalent shift was observed at 2 μM (Fig. 6B). Similar results were observed if the *fruA2* probe was instead used (data not shown). These results indicated that His_6_-MrpC_AAAA_ affinity for both the *mrpC* and *fruA* probes was reduced approximately 4-fold. Thus, replacing the TTSS motif in MrpC appears to reduce efficient binding of MrpC to promoters from which MrpC represses (*mrpC*) or induces (*fruA*) transcription. Interestingly, the His_6_-MrpC_ΔN25_ protein, which completely lacks the N-terminus binds with equal efficiency to *mrpC* (McLaughlin *et al*., 2018) and *fruA* (data not shown) binding sites.

Most Crp/Fnr family transcriptional regulators are typically dimers in solution and ligand binding induces a conformational change that increases affinity for binding to target sequences on the DNA. However, *E. coli* Fnr activity is regulated by dimer-to-monomer transition upon oxidation of an Fe-S cluster coordinated by cysteine residues. Fnr also contains an amino-terminal extension, and three out of the four cysteines that coordinate the Fe-S cluster are located near the TTSS motif in MrpC (Fig. S4). To examine whether MrpC activity was likewise controlled by monomer-to-dimer transition, and whether the TTSS motif could be involved in this transition, we subjected 8 μM purified His_6_-MrpC, His_6_-MrpC_ΔN25_ or His_6_-MrpC_AAAA_ to gel filtration analysis on a Superdex^TM^ 200 10/300 GL column compared to molecular mass standards. For each run, the absorbance at 280 nm (A_280_) versus retention volume was recorded and 0.8 mL fractions were collected. Fractions corresponding to A_280_ peaks were examined by SDS-PAGE for MrpC. We observed peak His_6_-MrpC, His_6_-MrpC_ΔN25_, or His_6_-MrpC_AAAA_ in fractions corresponding to the A_280_ peaks at 15.3, 15.5, and 15.2 mL retention volumes, respectively (Fig. 6C). No MrpC was detected in the large peaks at ∼10 mL retention volume which may have arisen due to light scattering. Analysis of the standard proteins indicated these retention volumes corresponded to molecular mass estimation of ∼50-60 kDa, most consistent with dimer formation of MrpC; the calculated molecular masses of His_6_-MrpC, His_6_-MrpC_ΔN25_, and His_6_-MrpC_AAAA_ proteins is 30.6, 28.2 and 30.7 kDa, respectively. Thus, neither the deletion of the N-terminal region, nor substitution of the TTSS motif prevented formation of MrpC dimers *in vitro*. These results suggested the perturbation of the TTSS motif subtly altered the conformation of MrpC resulting in reduced affinity for target promoters.

## DISCUSSION

MrpC is a Crp/Fnr transcriptional regulator that is essential for induction of aggregation and sporulation in the *M. xanthus* developmental program. A long-standing model suggested that the Pkn8/Pkn14 serine/threonine kinase cascade repressed MrpC activity by phosphorylating MrpC on threonine residues within its amino-terminal 25 residues (N25). Specifically, *in vitro* analyses showed that Pkn14 was sufficient to phosphorylate MrpC, and Pkn8 phosphorylates Pkn14, but not MrpC (Nariya & Inouye, 2005b). It was assumed that the threonine at position 21 and/or 22 was the phosphorylation target, because Pkn14 did not phosphorylate MrpC2, and thin layer chromatography of acid-hydrolyzed MrpC∼P indicated the phosphorylated residues corresponded to phospho-threonine. Phosphorylation was proposed to prevent proteolytic processing into MrpC2 (aka MrpC_ΔN25_), thought to be a more active isoform (Nariya & Inouye, 2006). This model was based on the observation that: 1) deletion of either *pkn8* or *pkn14* produced an early developmental phenotype, 2) ‘MrpC2’ was observed at higher levels in these strains, and 3) purified MrpC_ΔN25_ bound with higher affinity to target DNA sequences *in vitro*. However, we have recently demonstrated that the MrpC N25 is essential for activity *in vivo*, MrpC2 appears to be an artifact generated during cell lysis (McLaughlin *et al*., 2018), and the observation that ‘MrpC2’ binds to target DNA sequences with higher affinity has not been reproduced (Robinson *et al*., 2014, McLaughlin *et al*., 2018). Therefore, we set out to elucidate the functional consequences of threonine phosphorylation in the MrpC amino-terminal region by the Pkn8/Pkn14 serine/threonine kinase cascade. Focusing first on a putative TTSS phosphorylation motif in N25, we demonstrated that substitution to AAAA inactivated MrpC *in vivo* (Figs. 1A, 2, and S2). However, as Pkn14 did not phosphorylate the TTSS directly *in vitro* (Fig. 3B), we instead conclude that the TTSS motif is necessary for appropriate recognition of MrpC by Pkn14 (Fig. 3B) and efficient binding to target promoters (Fig. 6). We also reveal *M. xanthus* strain-specific activation of the Pkn8/14 kinase cascade (Fig. 4). Finally, our data reveal MrpC has an important role in stabilizing *M. xanthus* development likely against micro-environmental and/or intrinsic noise (Figs. 2 and S2). We propose a revised model for control of MrpC activity that reconciles our data with previous observations; this model is presented in two parts below.

### A revised model for Pkn8/Pkn14-dependent phosphorylation of MrpC

Our data revealed that Pkn8 and Pkn14 are not normally activated (i.e. auto-phosphorylated) under laboratory conditions in the wild type DZ2 *M. xanthus* strain, because kinase-dead versions of either (or both proteins) do not appreciably affect development (Fig. 4A and B). However, given the strong phenotype for both DZF1 *pkn14*_K48N_ (Nariya & Inouye, 2008) and DZF1 Δ*pkn14* (Nariya & Inouye, 2005b) mutations (Fig. S3), we speculate that in the DZF1 background, Pkn14 is activated (Pkn14∼P) resulting in a predominantly phosphorylated MrpC (MrpC∼P) species, that represses MrpC activity. Therefore, relative to the parent, the DZF1 *pkn14* mutant displays an obvious early aggregation phenotype (Nariya & Inouye, 2005b)(Fig. S3), because MrpC activity is no longer repressed by phosphorylation. The stimulus activating Pkn14 (perhaps via Pkn8) may be related to envelope stress or altered energy stores, because the DZF1 background contains the partially defective *pilQ1* allele encoding a major outer membrane secretin necessary for efficient type IV-pili production (Wall *et al*., 1999). *M. xanthus* type IV pili are necessary for social motility and are connected to production of surface polysaccharides (Yang *et al*., 2010, Black *et al*., 2017, Hu *et al*., 2016), suggesting multiple energy intensive processes are likely altered in this background (Hu *et al*., 2012).

Another puzzling observation was that although kinase-dead Pkn14 (Pkn14_K48N_) did not display an obvious developmental phenotype in the DZ2 background, deletion of *pkn14* resulted in delayed development (Fig. 4A). These results suggest a specific role for unphosphorylated Pkn14 in efficient induction of development. This role seems to be largely independent of MrpC, because we do not observe drastic changes in MrpC or FruA levels (targets of MrpC activity) in the Δ*pkn14* mutant (Fig. 4C). Pkn14 (along with Pkn8) belongs to a large kinase / scaffold protein network (Nariya & Inouye, 2005c), which includes at least two other kinases, Pkn9 and Pkn1, which appear to induce and repress aggregation, respectively (Hanlon *et al*., 1997, Munoz-Dorado *et al*., 1991). We suggest unphosphorylated Pkn14 affects aggregation indirectly through these proteins, whereas when Pkn14 is stimulated to autophosphorylate, it instead represses development by direct phosphorylation of MrpC. Thus, rather than functioning as a core component in activation of MrpC, we suggest Pkn14 (and likely Pkn8) function to modulate the developmental program in response to certain environmental conditions. How could MrpC activity be repressed by phosphorylation? We did not detect *in vitro* Pkn14-dependent phosphorylation of MrpC on threonine residues in the amino terminus as previously proposed, but rather on adjacent serine threonine sites within the cNMP domain (Ser_55_ Thr_56_) and on four sites within the DNA binding domain (Fig. 3B). We have not demonstrated the functionality of these residues, but the observation that MrpC becomes phosphorylated in the DNA binding region is consistent with observations that Pkn14-dependent phosphorylation of MrpC∼P has been shown to reduce binding to *mrpC* and *fruA* promoters *in vitro* (Nariya & Inouye, 2006). An analogy can be drawn to the *Bradyrhizobium japonicum* (Bj) FixK_2_ Crp/Fnr family member, which regulates genes required for microoxic, anoxic, and symbiotic growth during root nodule symbiosis with soybean plants (Nellen-Anthamatten *et al*., 1998).

BjFixK_2_ binds target sequences in the absence of known ligand and modification of a cysteine residue (Cys_183_) located directly adjacent to the helix-turn-helix motif in the DNA binding domain reduces DNA affinity (Mesa *et al*., 2009). Specifically, reactive oxygen species (ROS), which likely indicate conditions are not ideal for nodulation, convert the cysteine thiol to a bulky, negatively charged, sulfinic/sulfonic acid derivative which is proposed to sterically hinder the BjFixK_2_-DNA interaction and repulse the phosphate backbone of the DNA (Bonnet *et al*., 2013a). Intriguingly, one of the MrpC residues targeted by Pkn14, Thr_191_ (Fig. 3B), corresponds by sequence alignment and MrpC secondary structure prediction to BjFixK_2_ Cys_183_ (Fig. S4A). Addition of a bulky negative charged phosphoryl group to Thr_191_ can be predicted to likewise hinder MrpC interactions with the DNA backbone.

The function of the additional MrpC residues observed to be phosphorylated by Pkn14 *in vitro* is unknown. While the observation that there are so many sites is surprising and certainly remains to be verified *in vivo*, multisite phosphorylation of transcription factors is a common phenomenon in eukaryotes contributing to integrated control of the intensity of transcription factor activation (Holmberg *et al*., 2002). Consistently, *M. xanthus* encodes an unusually large number of eukaryotic-like serine/threonine protein kinases (Perez *et al*., 2008) that are likely organized into integrated signaling networks that coordinate multiple physiological responses (Nariya & Inouye, 2005c) (Nariya & Inouye, 2002).

### The role of the MrpC amino terminal extension

It was previously proposed that the MrpC amino-terminal extension (N25), which is not present in Crp homologs, must be removed for full MrpC activity. However, it is now clear that deletion of N25 in MrpC renders the protein inactive *in vivo* (McLaughlin *et al*., 2018). An *mrpC*_ΔN25_ mutant is unable to: 1) aggregate or sporulate efficiently, 2) regulate *mrpC* expression by negative autoregulation (i.e. *mrpC* expression was not repressed), and 3) to induce FruA (McLaughlin *et al*., 2018). Here, we additionally showed that MrpC_ΔN25_ was not subject to efficient proteolytic turnover (Fig. 5B). The recent observations that purified MrpC_ΔN25_ protein binds with equal affinity as full length MrpC to DNA target sequences *in vitro* (McLaughlin *et al*., 2018, Robinson *et al*., 2014), strongly suggests that the amino terminus is not required for DNA binding *per se*, but is required for additional contacts necessary to control transcription *in vivo*.

Our initial hypothesis at the start of this study was that phosphorylation of one or more residues in the TTSS motif modulates these proposed interactions. First, we demonstrated that complete replacement of TTSS with alanines produced similar, but slightly less extreme, results as the *mrpC*_ΔN25_ strain *in vivo*. The *mrpC*_AAAA_ strain failed to aggregate and sporulate efficiently (Fig. 1B and C), displayed a reduced *mrpC* negative autoregulation (i.e. *mrpC* expression was inefficiently repressed) (Fig. 5A), induced FruA in efficiently (Fig. 5C) and was unable to effectively turnover MrpC_AAAA_ (Fig. 5B). In contrast to MrpC_ΔN25_, however, purified MrpC_AAAA_ displayed significantly reduced affinity for *mrpC* and *fruA* DNA binding sites (Fig. 6A and B). Circular dichroism analysis suggested that the MrpC_AAAA_ mutation did not result in a drastically different secondary structure from the wild type (Fig. 3C). This observation reassured us that the reduction in DNA binding was not because the MrpC_AAAA_ was partially unfolded and raised the possibility that the TTSS to AAAA substitution may prevent the amino-terminus from assuming a configuration that stabilizes the DNA binding conformation of the protein. Finally, we did not detect Pkn14-dependent phosphorylation on the TTSS motif *in vitro* (Fig. 3B) and observed a general decrease in Pkn14-dependent phosphorylation on MrpC_AAAA_, suggesting inefficient recognition of MrpC_AAAA_ by Pkn14. We favor the interpretation that the entire amino terminus serves as a general protein interaction region, consistent with its intrinsic disorder prediction (data not shown). The TTSS motif in particular may act as a polar motif that allows stabilization of protein interactions, such as cooperative interactions with itself (Nariya & Inouye, 2006), contact with RNAP (Korner *et al*., 2003), FruA (Korner *et al*., 2003), or proteins that modulate the developmental program through MrpC, as we propose for Pkn14.

Many Crp/Fnr family members contain amino terminal extensions which are required for activity. For instance, the *E. coli* Fnr amino terminal extension contains three out of four of the cysteine residues which coordinate the Fe-S cluster that is necessary to sense anaerobic conditions (Spiro & Guest, 1990)(Fig. S4A). Intriguingly, MrpC structure predictions (Kelley *et al*., 2015) model the TTSS motif at the end of an alpha helix that threads between the dimerization helix and the β-sheet scaffold and near to one of DNA binding helices in the helix-turn-helix motif (Fig. S5A). This arrangement has been seen in the crystal structure of *Mycobacterium tuberculosis* (Mt) Cmr (Ranganathan *et al*., 2018), where the amino terminal extension (helix N1) is predicted to play a role in modulating DNA binding and/or dimerization perhaps in response to cellular signals (see Fig. S4A for alignments). Thus, interaction with other proteins, or unknown ligands, may reorient the amino-terminus. The TTSS motif may play a role in modulating this reorientation. Our on-going efforts to solve the MrpC structure may illuminate how the region could affect MrpC activity.

Interestingly, MrpC shares many architectural features with the Fnr-like branch of transcriptional regulators that regulate gene transcription in response to anaerobic or microoxic conditions. Like MrpC, BjFixK_2_, as well as *E. coli* (Ec) Fnr, are regulated by proteolytic turnover (Bonnet *et al*., 2013b, Mettert & Kiley, 2005). BjFixK_2_ contains proteolytic sequence determinants in the extreme C-terminus of the protein which are recognized and degraded by ClpAP_1_ (Bonnet *et al*., 2013b)(Fig. S4). In the case of EcFnr, sequence determinants are found in the both the amino terminal 5-11 residues and in the last two residues of the protein (residues 249 and 250)(Fig. S4), and proteolytic turnover is dependent on ClpXP (Mettert & Kiley, 2005). We have shown here that both MrpC_AAAA_ and MrpC_• N25_ are not subject to proteolysis (Fig. 5B) suggesting that the amino terminus may contain the sequence determinants for proteolytic turnover. However, this interpretation is complicated by the observation that EspA is highly reduced in the *mrpC*_AAAA_ background (data not shown). EspA is an integral component of the signaling system that induces proteolytic turnover of MrpC (Mettert & Kiley, 2005), and *espA* expression is dependent on MrpC which binds to the *espA* promoter *in vitro* (A. Schramm and P. Higgs, unpublished results).

### MrpC plays a role in stabilizing the developmental program

A surprising finding from this study is that TTSS motif mutants bearing non-consecutive T/S residues produce highly variable phenotypes ranging from wild type to early, delayed, or retro development in different clones and in different biological replicates of the same clone (Figs. 2 and S2). These results reveal a previously unrecognized role for MrpC in maintaining developmental stability. It has been argued that developmental systems have evolved mechanisms to buffer against noisiness due to stochastic variation in numbers of regulatory molecules that must interact to promote progression through developmental pathways (Nijhout & Davidowitz, 2003, DeLaurier *et al*., 2014). In the case of MrpC TTSS motif variants and *M. xanthus* development, this process most likely involves micro-environmental canalization, or buffering against phenotypic variation due to fine-scale environmental variation (such as slight differences in nutrient concentration or cell density between assays) and developmental noise (Nijhout & Davidowitz, 2003). Consistently, it has been recently observed that ultra-sensitive responses to nutrient concentration are mediated by MrpC (Hoang & Kroos, 2018).

MrpC’s role as an environmental capacitor is likely the result of its position as a hub in the genetic regulatory network. Multiple signaling systems feed into MrpC (Schramm *et al*., 2012, Higgs *et al*., 2008, Stein *et al*., 2006, Nariya & Inouye, 2005a, Inouye & Nariya, 2008, Hoang & Kroos, 2018, Rajagopalan & Kroos, 2017), and MrpC directly and indirectly induces regulatory feedback loops (Kroos, 2007) (Hoang & Kroos, 2018). Furthermore, MrpC accumulation correlates with distinct cell fates: little or no MrpC is found in peripheral rods and highest accumulation in fruiting body cells (Lee *et al*., 2012) and misaccumulation of MrpC can lead to inappropriate cell fate segregation (Cho & Zusman, 1999, Schramm *et al*., 2012). Uncoordinated and/or inefficient MrpC interactions with target promoters or binding partners, or misaccumulation in inappropriate cell types, could result in stochastic phenotypes. Consistent with the observation that MrpC may be phosphorylated on multiple sites, modeling approaches have suggested that multisite phosphorylation is mechanism to filter noise (Aledo, 2018). We are currently examining whether MrpC is phosphorylated *in vivo* in response to changing environmental conditions, whether the amino terminus is a general interaction motif, and whether additional STPKs play a role in this process.

## Experimental Procedures

### Bacterial growth and development conditions

*E. coli* cells were grown at 37 °C on LB (0.1% tryptone, 0.5% yeast extract, 0.5% NaCl) in agar plates (1.5%) or with shaking (220 rpm) in LB broth supplemented with 50 µg ml^-1^ kanamycin, 100 µg ml^-1^ ampicillin, and/or 34 µg ml^-1^ chloramphenicol as needed. Vegetative *M. xanthus* cells were grown at 32 °C on CYE (0.1% casitone, 0.5% yeast extract, 10 mM MOPS-KOH, pH 7.6, 8 mM MgSO_4_) 1.5% agar (Campos & Zusman, 1975) supplemented with 100 µg ml^-1^ kanamycin when necessary, or in CYE broth (CYE without agar) with shaking at 220 rpm.

*M. xanthus* strains were induced to develop under submerged culture (Lee *et al*., 2010) unless otherwise indicated. Briefly, cells were grown overnight in CYE broth, diluted to 0.035 A_550_ in CYE and 0.5, 8, or 16 mL of cells was seeded into 24 well tissue culture plates, or 60 mm or 150 mm petri dishes, respectively. Cells were grown into a confluent layer at 32°C for 24 hours, then CYE was replaced with an equivalent volume of MMC (10 mM MOPS pH 7.6, 4 mM MgSO_4_, 2 mM CaCl_2_) to induce development. Cells were incubated undisturbed at 32°C, and pictures were recorded with a Leica DMC 2900 stereo microscope.

For high-throughput, high resolution development imaging (Glaser & Higgs, 2019), cells were induced to develop by submerged culture as described above, except 0.15 mL cells were seeded into each well of a 96 well microtiter plate and incubated at 32 °C. After 24 hours of development, plates were transferred to a Tecan Infinite M200 plate reader (pre-warmed to 32 °C), and images were recorded from each well every 30 min from 24-72 hours post starvation using the plate reader cell confluence feature. Images were then assembled into movies in ImageJ (Schneider *et al*., 2012). For each movie, the time frame of developmental stages (onset of aggregation, initial aggregates, aggregates after consolidation/dissolution, mature aggregates, and darkened fruiting bodies) were manually recorded.

To determine the number of spores produced during development, cells were induced to develop by submerged culture in 24 well plates as described above. After 72 or 120 hours of starvation, cells from triplicate wells were harvested, pelleted at 17,000 x g for 5 min, supernatant was removed, and pellets stored at -20°C or used immediately. To kill non-spores, pellets were resuspended in 0.5 mL of water, heated at 50°C for 60 min, and sonicated three times for 30 sec (0.5 sec on 0.5 sec off) at 30% output on Branson Sonifier 250a equipped with a microtip. The remaining spherical and phase bright spores were enumerated on a Hawksley Helber bacteria hemocytometer.

The number of chemically-induced spores were determined as described previously (Holkenbrink *et al*., 2014). Briefly, 25 mL cultures *M. xanthus* cells were grown at 32°C in CYE broth to an OD of ∼ 0.3 A_550_, 10 M sterile glycerol was added to a final concentration of 0.5 M and cultures were incubated at 32°C for 24 hours. The culture was pelleted and resuspended in 5 mL sterile water, and 3 x 0.5 mL were heated at 50°C for one hour and sonicated and counted as described above. Spore number was reported as percent of starting cells, where an OD of 0.7 A_550_ corresponded to 4×10^8^ cells mL^-1^

### Construction of plasmids and strains

Plasmids used to construct *M. xanthus* strains (Table 1) were constructed using standard restriction enzyme/ligation cloning techniques followed by transformation into *E. coli* strain TOP10. For plasmids used to generate strains bearing in-frame deletions or point mutations in the endogenous *M. xanthus* locus, gene fragments contained ∼500 bp fragments upstream and downstream from the desired deletion or point mutation and were constructed by over-lap PCR as described previously in detail (Lee *et al*., 2010), and cloned into pBJ114 using the primers and enzymes listed in Table S2. For plasmids used to express genes at the exogenous Mx8 phage attachment site, pFM18 (kan^R^, Mx8 *attP*) was used. Plasmid insert fragments containing the native promoter and desired mutated gene were initially constructed by overlap PCR or by direct PCR amplification using the templates, primers, and enzymes listed in Table S2. For plasmids used to express genes at the 1.38 kb *M. xanthus* genome integration site, P_mrpC_-*mrpC* inserts were cloned into pMR3679 (km^R^, 1.38 kb recombination fragment), such that the vanillate promoter was removed. Plasmid inserts were constructed as indicated in Table S2.

**Table 1.** Strains and plasmids used in this work.

| ***M. xanthus* strains** |  |  |
| --- | --- | --- |
| **Strain** | **Genotype** | **Source** |
| DZ2 | Wild type | (Campos & Zusman, 1975) |
| PH1025 | DZ2 $\Delta mrpC$ | (Lee *et al.*, 2012) |
| PH1108 | DZ2 $mrpC_{\Delta N25}$ | (McLaughlin *et al.*, 2018) |
| PH1139 | DZ2 $mrpC_{AAAA}$ | This study |
| PH1118 | PH1025 $attB::P_{mrpC}-mrpC$, Km<sup>R</sup> | (McLaughlin *et al.*, 2018) |
| PH1530 | PH1025 $attB::P_{mrpC}-mrpC_{AAAA}$, Km<sup>R</sup> | This study |
| PH1531 | PH1025 $attB::P_{mrpC}-mrpC_{TAAA}$, Km<sup>R</sup> | This study |
| PH1532 | PH1025 $attB::P_{mrpC}-mrpC_{AAAS}$, Km<sup>R</sup> | This study |
| PH1547 | PH1025 $attB::P_{mrpC}-mrpC_{TASA}$, Km<sup>R</sup> | This study |
| PH1533 | PH1025 $attB::P_{mrpC}-mrpC_{ETSS}$, Km<sup>R</sup> | This study |
| PH1132 | DZ2 $\Delta pkn14$ | This study |
| PH1133 | DZ2 $pkn14_{K48N}$ | This study |
| PH1347 | DZ2 $\Delta pkn8$ | This study |
| PH1548 | DZ2 $pkn8_{K116N}$ | This study |
| PH1549 | PH1133 $pkn8_{K116N}$ | This study |
| PH1100 | DZ2 $attB::P_{mrpC}-mCherry$, Km<sup>R</sup> | (McLaughlin *et al.*, 2018) |
| PH1104 | PH1025 $attB::P_{mrpC}-mCherry$, Km<sup>R</sup> | (McLaughlin *et al.*, 2018) |
| PH1306 | PH1139 $attB::P_{mrpC}-mCherry$, Km<sup>R</sup> | This study |
| ***E. coli* strains** |  |  |
| **Strain** | **Genotype** | **Source** |
| TOP10 | F<sup>-</sup> *endA1 recA1 galE15 galk16 nupG rpsL*<br>$\Delta lacX74 \Phi 80 lacZ \Delta M15 araD139 \Delta(ara, leu)7697$<br>*mcrA* $\Delta(mrr-hsdRMS-mcrBC)$ $\lambda^{-}$ | Invitrogen |
| BL21 λDE3 | F<sup>-</sup> *ompT gal dcm lon hsdS<sub>B</sub> (r<sub>B</sub><sup>-</sup> m<sub>B</sub><sup>-</sup>)* $\lambda$ (DE3 [*lacI* *lacUV5-T7* gene 1 *ind1 sam7 nin5*]) | Novagen |
| **Plasmid** | **Genotype** | **Source** |
| **Mutagenesis plasmids** |  |  |
| pBJ114 | Suicide plasmid with Km<sup>R</sup> and galk | (Julien *et al.*, 2000) |
| pVG129 | pBJ114 $mrpC_{AAAA}$ | This study |
| pFM18 | Mx8 phage *attP* integration plasmid | (Muller *et al.*, 2010) |
| pBEF010 | pFM18 $P_{mrpC}-mrpC_{AAAA}$ | This study |
| pBEF007 | pFM18 $P_{mrpC}-mrpC_{TAAA}$ | This study |
| pBEF011 | pFM18 $P_{mrpC}-mrpC_{AAAS}$ | This study |
| pBEF015 | pFM18 $P_{mrpC}-mrpC_{TASA}$ | This study |
| pBEF006 | pFM18 $P_{mrpC}-mrpC_{ETSS}$ | This study |
| pXM001 | pBJ114 $\Delta pkn14$ | This study |
| pVG125 | pBJ114 *pkn14*<sub>K48N</sub> | This study |
| pXM002 | pBJ114 Δ*pkn8* | This study |
| pXM003 | pBJ114 *pkn8*<sub>K116N</sub> | This study |
| pPTM014 | Mx8 *attP*, P<sub>*mrpC*</sub>-*mCherry* | (McLaughlin *et al.*, 2018) |
| **Overexpression plasmids** |  |  |
| pET28a+ | T7 promotor, His<sub>6</sub> tag (N-terminal), *Km*<sup>R</sup> | Novagen |
| pPH158 | pET28a+ *mrpC*, *Km*<sup>R</sup> | (Lee *et al.*, 2012) |
| pPH167 | pET28a+ *mrpC*<sub>ΔN25</sub>, *Km*<sup>R</sup> | (McLaughlin *et al.*, 2018) |
| pVG131 | pET28a+ *mrpC*<sub>AAAA</sub>, *Km*<sup>R</sup> | This study |
| pASK-IBA15+ | *tet* promotor, *Strep* tag II (N-terminal), *Ap*<sup>R</sup> | IBA Lifesciences |
| pPH166 | pASK-IBA15+ *pkn14* | This study |
| pVG130 | pASK-IBA15+ *pkn14*<sub>K48N</sub> | This study |
| pET32a+ | T7 promotor, Trx, His<sub>6</sub> tag, *Ap*<sup>R</sup> | Novagen |

Strains bearing point mutations or in-frame deletions in the endogenous genetic locus (Tables 1 and S1) were constructed using the strategy described in detail previously (Lee *et al*., 2010), Briefly, pBJ114 (*km*^R^, *galK*) based plasmids were introduced into the relevant *M. xanthus* strains by electroporation, and plasmid integration through homologous recombination was selected by kanamycin-resistance. Plasmid loop-out via a second homologous recombination event was screened by growth on 2-5% galactose, and clones with the desired mutation were screened by PCR (in-frame deletions) or PCR amplified and then sequenced (point mutations) (Lee *et al*., 2010).

To generate strains bearing genes expressed from the Mx8 phage attachment site (*attB*), pFM18 based plasmids (Tables 1 and S1) were electroporated into the relevant *M. xanthus* strains and site-specific recombination into *attB* selected by kanamycin resistance and screened for correct integration by PCR as described previously (McLaughlin *et al*., 2018). For strains bearing plasmid integrations at the 1.38kb integration site, pMR3679-based plasmids were electroporated into the relevant *M. xanthus* strains and homologous recombination into the 1.38 kb *M. xanthus* genome region was selected by kanamycin resistance. Resulting colonies were screened by PCR for correct integration as described previously (Garcia-Moreno *et al*., 2009). For all constructed strains, the developmental phenotypes of at least three independent clones were assayed to confirm a true phenotype.

Protein expression plasmids were constructed using the templates, oligonucleotides, and enzymes listed in Table S2.

### Immunoblot analysis

To generate cell lysates for immunoblot analysis, *M. xanthus* was induced to develop in 150 mm petri dishes, as described above. At the indicated time points, cells were harvested, pelleted at 4696 x g at 4°C, and cell pellets were either used immediately or stored at -20°C. Cell pellets were resuspended in 0.5 mL of cold MMC, 0.5 mL of cold, 26% TCA was added, and tubes were incubated on ice for at least 15 minutes.

Precipitated proteins were pelleted at 4°C at 17,000 x g for 5 minutes, supernatant was removed, the pellet was resuspended in 100% ice-cold acetone, centrifuged for 1 minute at room temperature, and repeated. After the second acetone wash, protein pellets were resuspended in 100 µL of 100 mM Tris pH 8.0, 300 µL of clear LSB (62.5 mM Tris-HCl, pH 6.8, 10% (v/v) glycerol, 2% (w/v) SDS) was added, and samples were heated at 94°C for 1 minute. Protein concentration in each sample was determined using a BCA Protein Assay Kit (Pierce). Samples were diluted to 1 µg µL^-1^ in 2X LSB (125 mM Tris-HCl, pH 6.8, 20% (v/v) glycerol, 4% (w/v) SDS, 10% (v/v) 2-mercaptoethanol, 0.02% (w/v) bromophenol blue), heated at 99°C for 5 minutes, and then stored at -20 °C. Protein samples were resolved by sodium dodecyl sulfate polyacrylamide gel electrophoresis (SDS-PAGE) on a 13% (MrpC and FruA), or 9 % (Pkn14) polyacrylamide gel, then transferred to a polyvinylidene fluoride (PVDF) membrane (Millipore) in a semi-dry transfer apparatus (Hoefer TE77X). For Western blots, membranes were either blocked for one hour at room temperature or overnight at 4°C in PBS (8 mM Na_2_HPO_4_, 2 mM KH_2_PO_4_ pH 7.4, 135 mM NaCl, 3.5 mM KCl) supplemented with 0.1 % Tween and 5 % non-fat milk. Primary antibodies were used at 1:500 (anti-MrpC) (Garcia-Moreno *et al*., 2009), 1:1000 (anti-FruA) (Lobedanz & Sogaard-Andersen, 2003), or 1:500 (anti-Pkn14). Goat anti-rabbit, HRP-conjugated secondary IgG secondary antibodies were used at 1:20 000 (Pierce).

Chemiluminescence (ECL Western Blotting Substrate, Pierce) and autoradiography film (Andwin Scientific) were used for detection of immune complexes. For quantitation, autoradiographs were scanned, and ImageQuant TL (GE Healthcare Life Sciences) was used to quantify background-subtracted signal intensities.

Polyclonal rabbit anti-Pkn14 antibodies were generated by Eurogentec (Serain, Belgium) using soluble purified Strep-Pkn14 protein. Purification was performed as described below. Anti-Pkn14 antibodies were purified from serum against Step-Pkn14 protein as per (Jagadeesan *et al*., 2009).

### Overexpression and purification of recombinant proteins

Overexpression plasmids (Table 2) for recombinant His_6_-tagged -MrpC, - MrpC_ΔN25_, - MrpC_AAAA_, or Trx-His_6_, were freshly transformed into *E. coli* BL21(λDE3) cells and ∼ 3 resulting colonies were grown overnight in 5 ml starter cultures, and then subcultured (1:100) into 500 ml LB broth containing the appropriate antibiotic. His-tagged proteins were induced in mid-log cells with 0.5 to 1 mM isopropyl 1-thio-β-D-galactopyranoside (IPTG) for 3 hours at 25 or 37 °C. His-tagged proteins were then purified at 4°C either by gravity flow (phosphotransfer and EMSA analyses) or FPLC (circular dichroism and gel filtration analyses) using Ni-NTA resin (Qiagen) or 5-mL HisTrap column (GE Healthcare), respectively. For purification of His_6_-tagged proteins by gravity flow, cell pellets were resuspended in 12 mL Lysis Buffer (50 mM HEPES-NaOH pH 7.4, 500 mM NaCl, 20 mM imidizole) containing 1 mg mL^-1^ lysozyme and 1:100 protease inhibitor cocktail (Sigma), and lysed by French press (18,000 psi, 3 times). Unlysed cells and/or inclusion bodies were removed by centrifugation at 600 x g at 4°C for 30 min and supernatant was applied to 0.5 mL Ni-NTA resin pre-equilibrated with lysis buffer, washed with 5 column volumes (CV) Wash Buffer (50 mM HEPES pH 7.4, 500 mM NaCl, 20 mM imidazole), and step-eluted with 3CV Elution Buffer (50 mM HEPES pH 7.4, 500 mM NaCl) containing 100 mM, 250 mM and then 500 mM imidazole. Elution fractions containing peak amounts of purified protein were pooled, dialyzed in 900 ml Dialysis Buffer 1 (50 mM HEPES pH 7.5, 250 mM NaCl, 1 mM dithiothreitol (DTT), 10% (v/v) glycerol) for 1 hour at room temperature, then in 900 ml dialysis buffer 2 (Dialysis Buffer 1 except 100mM NaCl and 20% glycerol) overnight at 4°C. Dialyzed proteins were stored at -20°C. For FPLC purification of His-tagged proteins, induced cells were resuspended in buffer B (20 mM Tris-HCl, 500 mM NaCl, 20 mM Imidazole pH 8.5) and lysed by passing the cell suspension 3-5 times through an EmulsiFlex-C3 homogenizer at 5,000-10,000 psi. Clarified lysate was loaded onto a 5-mL HisTrap column (GE Healthcare) and weakly bound proteins were eluted off with a linear gradient from 20-300 mM imidazole (buffer B to buffer C) followed by elution of His-tagged MrpC proteins with buffer C (20 mM Tris-HCl, 100 mM NaCl, 900 mM Imidazole pH 8.9). Fractions containing MrpC were pooled, concentrated, buffer-exchanged in buffer B, and stored at 4 °C until further use.

Overexpression plasmids (Tables 1 and S2) for recombinant Step-tagged –Pkn14, or – Pkn14_K48N_, were freshly transformed and induced for overexpression as described above except that the inducing agent was 0.06 µg mL^-1^ of anhydrotetracycline for 3 hours at 20°C. Cultures were then pelleted at 15,000 rpm for 15 minutes at 4°C, and cell pellets were stored at -20°C until needed. For purification of Strep-tagged proteins, cell pellets were resuspended in 30mL Buffer W (100 mM Tris pH 8.0, 150 mM NaCl, 1 mM EDTA) containing 1mg mL^-1^ lysozyme and 1:100 protease inhibitors, lysed by French press, and cleared as described above. Cell supernatants were loaded onto columns containing 0.8 ml Strep-Tactin resin (IBA) pre-equilibrated with Buffer W. Columns were washed with 5 CV Buffer W and Strep-tagged proteins were eluted with 4 CV Buffer E (100 mM Tris pH 8.0, 150 mM NaCl, 1 mM EDTA, 2.5 mM desthiobiotin); 0.8 mL fractions were collected, analyzed by SDS-PAGE, and elution fractions containing peak amounts of purified protein were pooled, dialyzed twice as described above except dialysis buffer contained 100 mM Tris pH 8.0, 150 mM NaCl, 1 mM DTT, 20% glycerol above, and stored at -20°C.

### In vitro kinase reactions and analysis

Kinase reactions were carried out by incubating 6 µg of each purified kinase and substrate in buffer P (50 mM Tris pH 8.0, 2.4 mM ATP, 6 mM MgCl_2_, 1 mM DTT) at 30°C for 30 minutes (typically in 30µL total volume), and quenched with an equal volume of 2 X LSB. 18 µL of each reaction was analyzed by SDS-13% PAGE. 1µL of phosphoprotein Molecular Weight Standards (Invitrogen) was included on the gel.

Protein phosphorylation was visualized by incubating the gel in ProQ Diamond® phosphoprotein stain (Invitrogen) according to the manufacturer’s instructions. Briefly, gels were incubated in 100 mL ProQ Diamond® fixing solution for at least 30 minutes, washed twice with water, then stained in ProQ Diamond® phosphoprotein stain for at least 75 minutes in the dark. Gels were destained in ProQ Diamond® destain solution three times for 30 minutes, then washed in water prior to imaging on a Typhoon FLA 7000 with an excitation wavelength of 532 nm and an R670 emission filter. Total protein was visualized by subsequently incubating rinsed gels in 60 mL Sypro Ruby stain (Invitrogen) overnight, followed by washes in Sypro Ruby wash solution for 30 minutes, 100 mL water two times for 5 minutes and then imaged at 532 nm excitation wavelength and an O580 emission filter. Signal intensities were quantified using ImageQuant TL, and background-subtracted phosphoprotein signal was normalized to background-subtracted total protein signal, then plotted as the ratio of kinase active over kinase dead reactions.

### Mass spectrometry

For mass spectrometry analysis, Pkn14 kinase reactions with MrpC substrate were set up as described above, except that 7.5 µg of each protein was used and reactions were stopped by flash freeze in dry ice. Protein samples were then reduced with 5 mM dithiothreitol (30 min at 65LC) and alkylated with 20 mM iodoacetamide (30 min at room temperature) in 50 mM ammonium bicarbonate, and digested with 1:50 dilution of trypsin in 25 mM ammonium bicarbonate and 10% acetonitrile (overnight at 37LC). Peptides were dried and resuspended in a 2% trifluoroacetic acid and 65% acetonitrile solution that was saturated with glutamic acid. Phosphopeptide enrichment was performed using an AssayMAP Bravo automated liquid handler paired with TiO_2_ cartridges (Agilent Technologies) according to manufacturer’s instructions. Peptides were eluted with a 5% ammonium hydroxide solution, dried, and resuspended in 0.1% formic acid, 0.005% trifluoroacetic acid and 5% acetonitrile prior to LC/MS analysis.

Peptides were separated by UHPLC reverse phase chromatography using C18 columns and an EASY-nLC system (Thermo) before introduction into an Orbitrap Fusion mass spectrometer (Thermo). Settings for MS1 included a scan range of 375-1600 m/z at 240K resolution. For MS2 scans detected in the ion trap, peptides with +2 and +3 charges were fragmented by collision induced dissociation (CID) and peptides with charges +3 to +7 were fragmented by electron transfer dissociation (ETD). RAW files were searched with the Sequest algorithm within Proteome Discoverer (Thermo; ver 2.1) using 100 PPM and 0.6 Da tolerances for parent and fragment ions, respectively; fixed modification of carbamidomethylation of Cys; variable modifications of deamidation of Asn/Gln, oxidation of Met, and phosphorylation of Ser/Thr/Tyr; and up to 2 missed tryptic cleavages. Forward and reverse database searches were performed against a protein database for *Myxococcus xanthus* (downloaded from Uniprot on 2018- 04-19; 8101 sequences). Results were imported into Scaffold (Proteome Software; ver 4.8) and a subset database was searched using X! Tandem. Peptide identifications were thresholded at a 1% false discovery rate (FDR).

### Circular Dichroism

The CD-spectra of 56 μg ml^-1^ His_6_-MrpC or His_6_-MrpC_AAAA_ in 20 mM Tris-HCl pH 8 and 100 mM NaCl were determined immediately after spinning the protein for 15 minutes at 20,000 x g with a Jasco 815 CD spectrometer with a 1 mm pathlength in the range λ = 190-260 nm. The spectra were collected in one pass with a bandwidth of 1 nm, scanning speed of 100 nm/min, and response time of 2 seconds at room temperature, following (Greenfield, 2006).

### Analysis of mCherry fluorescence

Strains bearing the *mrpC* mCherry expression reporter (*attB*::P*_mrpC-_mCherry*) were analyzed for mCherry fluorescence as described previously (McLaughlin *et al*., 2018), except fluorescence was recorded in a Tecan Infinite M200 plate reader at 582 nm excitation and 613 nm emission wavelengths. Briefly, strains were induced to develop under submerged culture in 24 well plates, triplicate wells were harvested, cells were dispersed, and triplicate 150 µL samples were analyzed for mCherry fluorescence intensity and normalized to total protein detected from remaining samples at each time point. Average data from three independent biological replicates was plotted.

### Chloramphenicol shutoff assays

MrpC turnover was analyzed as described previously (Schramm *et al*., 2012). Briefly, strains were induced to develop by submerged culture in 150 mm petri dishes, as described above. Chloramphenicol was added to a final concentration of 34 µg mL^-1^ to one petri dish per time point for each strain, and cells were harvested at 0, 10, 20, 30, or 60 minutes after addition of chloramphenicol, pelleted at 4 696 x g at 4°C, resuspended in 400 µL of 70°C 2X LSB, then heated at 99°C for 5 minutes. Equal proportions of samples were subject to anti-MrpC immunoblot as described above. ImageQuant TL was used to quantify background-subtracted signal intensities from duplicate replicates, and the signal intensity for each time point was normalized to that of t = 0 for each strain. The natural log of those values was plotted against time, and the slope (k) of the linear fit was used to calculate the half-life (t_1/2_) of MrpC, using the equation t_1/2_ = ln(2)/-k

### Electrophoretic mobility shift assays

For electrophoretic mobility shift assays (EMSAs) were performed as described in detail, previously (McLaughlin *et al*., 2018). Briefly, DNA probes consisted of three annealed primers: a double Cy-5 labeled universal forward primer, an unlabeled forward primer that contained the protein binding site, and an unlabeled reverse primer that was complementary to both forward primers that were annealed in a thermocycler. The indicated concentrations of purified His_6_-MrpC or His_6_-MrpC_AAAA_ were incubated with 50 nM labeled probe for 30 min at 20°C and then resolved on 10% polyacrylamide (37.5:1 acrylamide/bis-acrylamide gels in 0.5x TBE at 100V at 4°C. Gels were imaged with a Typhoon FLA 7000 [Pixel size: 25 μm, PMT: 500, Latitude: 4, Excitation wavelength: 635 nm, Emission filter: R670].

### Gel filtration analyses

Analytical size-exclusion chromatography was carried out on a Superdex 200 10/300 GL column (GE Healthcare) equilibrated in 20 mM Tris-HCl, 500 mM NaCl, 20 mM imidazole pH 8.5 by an ÄKTA High Performance Liquid Chromatography (HPLC) apparatus. Eight μM His_6_-MrpC, - MrpC_ΔN25_ or -MrpCAAAA proteins were applied with a flow rate of 0.5 ml min^-1^. 0.8 mL fractions were collected, precipitated with trichloroacetic acid (TCA) and analyzed by SDS-PAGE. Molecular weight standards were used to calibrate the column: alcohol dehydrogenase (150 kDa), albumin (66 kDa), carbonic anhydrase (29 kDa), and myoglobin (17.6 kDa) had retention volumes of 13.0 mL, 14.67 mL 16.75 mL, and 17.35 mL respectively.

## Acknowledgements

We gratefully acknowledge Xiaowei Mei for initial construction of *pkn8* and *pkn14* plasmids, Brian Nguyen for MrpC purification assistance, Petra Mann for technical assistance, and Juan A. Arias Del Angel for stimulating discussions. This work was funded by the National Science Foundation Career award IOS-1651921, Wayne State startup funds, Wayne State Chemistry Biology Interface (CBI) fellowship (BF) and the Max Planck Society (PH and VB).

## Author Contributions

PIH conceived of the study. BF, VB, PTM, SD, GB and PIH acquired and/or analyzed data. PIH and BF wrote the manuscript.

## Supporting Information

**Table S1.** Strains and plasmids used in supplementary data.

**Table S2.** Oligonucleotides and construction of plasmids used in this study.

**Figure S1.** Developmental phenotypes of MrpC TTSS motif mutants.

**Figure S2.** Insertion of P*_mrpC_*-*mrpC* alleles at the 1.38kb *M. xanthus* genomic site does not rescue loss of developmental robustness.

**Figure S3.** Developmental phenotype of Δ*pkn14* in the DZF1 versus DZ2 backgrounds.

**Figure S4.** Alignments of MrpC with Crp/Fnr homologs reveals functional residue conservation.

**Figure S5.** Predicted MrpC tertiary structure.

