## Supplemental Data for "An amino-terminal threonine/serine motif is necessary for activity of the Crp/Fnr homolog, MrpC, and for *Myxococcus xanthus* developmental robustness"

**SUPPLEMENTAL FIGURES**

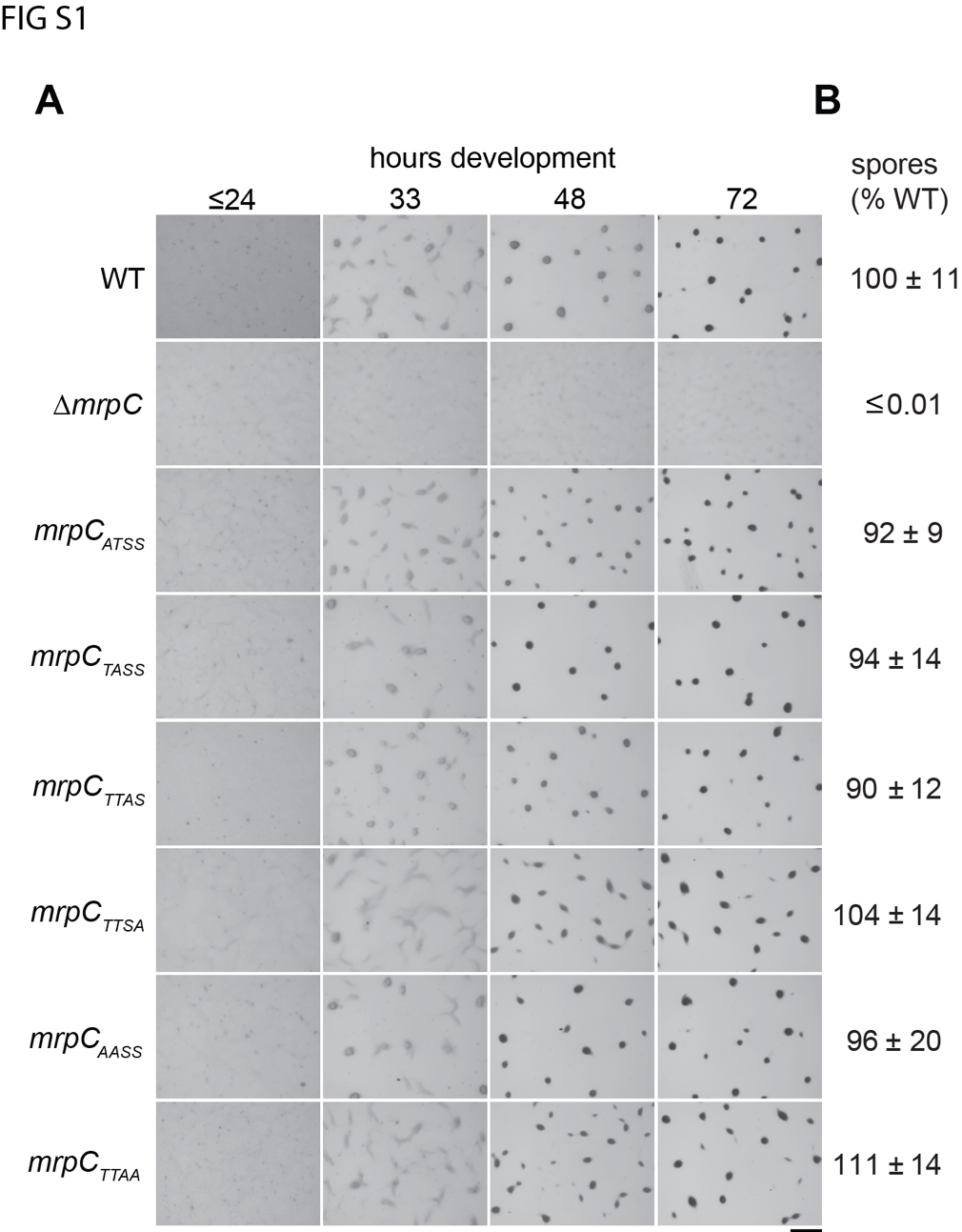

Figure S1. Developmental phenotypes of MrpC TTSS motif mutants. A. Wild type (wt: DZ2), Δ*mrpC* (PH1025), *mrpC_ATSS_* (PH1154), *mrpC_TASS_* (PH1155), *mrpC_TTAS_* (PH1137), *mrpC_TTSA_* (PH1157)¸ *mrpC_AASS_* (PH1136), *mrpC_TTAA_* (PH1138), strains were induced to develop under submerged culture and pictures recorded at the indicated hours of development. Bar: 0.5 mm. B. Percent of wild type heat and sonication resistant spores harvested from cells developed for 72 hours under submerged culture. Values are the average and associated standard deviations from three independent biological replicates.

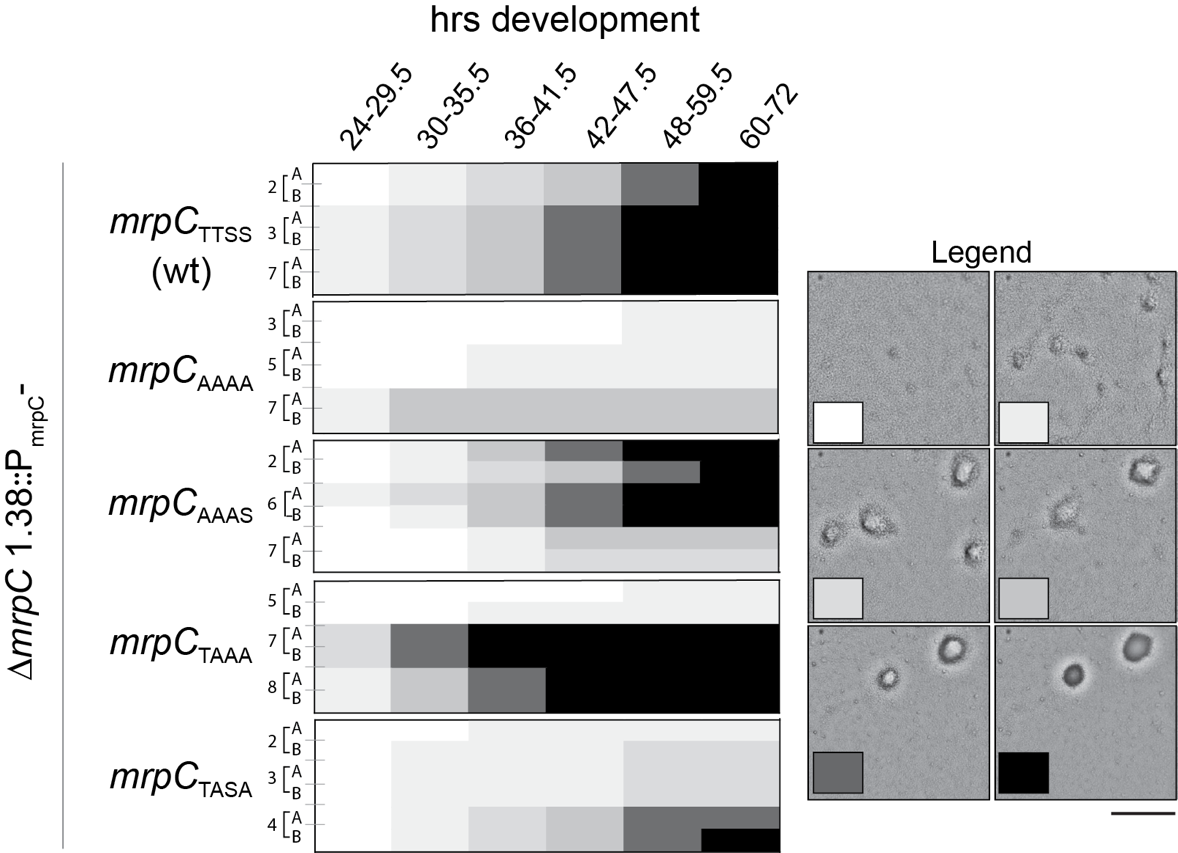

Figure S2. Insertion of P*_mrpC_*-*mrpC* alleles at the 1.38kb *M. xanthus* genomic site does not rescue loss of developmental robustness. Inter- and intra-clone variability in the developmental phenotypes of *mrpC*_TTSS_ (wild type; PH1539), *mrpC_AAAA_* (PH1540)*, mrpC_AAAS_* (PH1542), *mrpC_TAAA_* (PH1541)*,* and *mrpC_TASA_* (PH1543) genes integrated at the 1.38kb site (Garcia-Moreno *et al.*, 2010) in the Δ*mrpC* background. The heat maps shown indicate the variability in the timing and extent of completion of fruiting body formation during development under submerged culture in 96 well plates in the indicated ranges of developmental times. Replicates are indicated by letters (A, B), and independent clones are represented by clone number. The shade of box corresponds to the extent of development as indicated by representative pictures in the legend (see text for details). Bar: 250 µm.

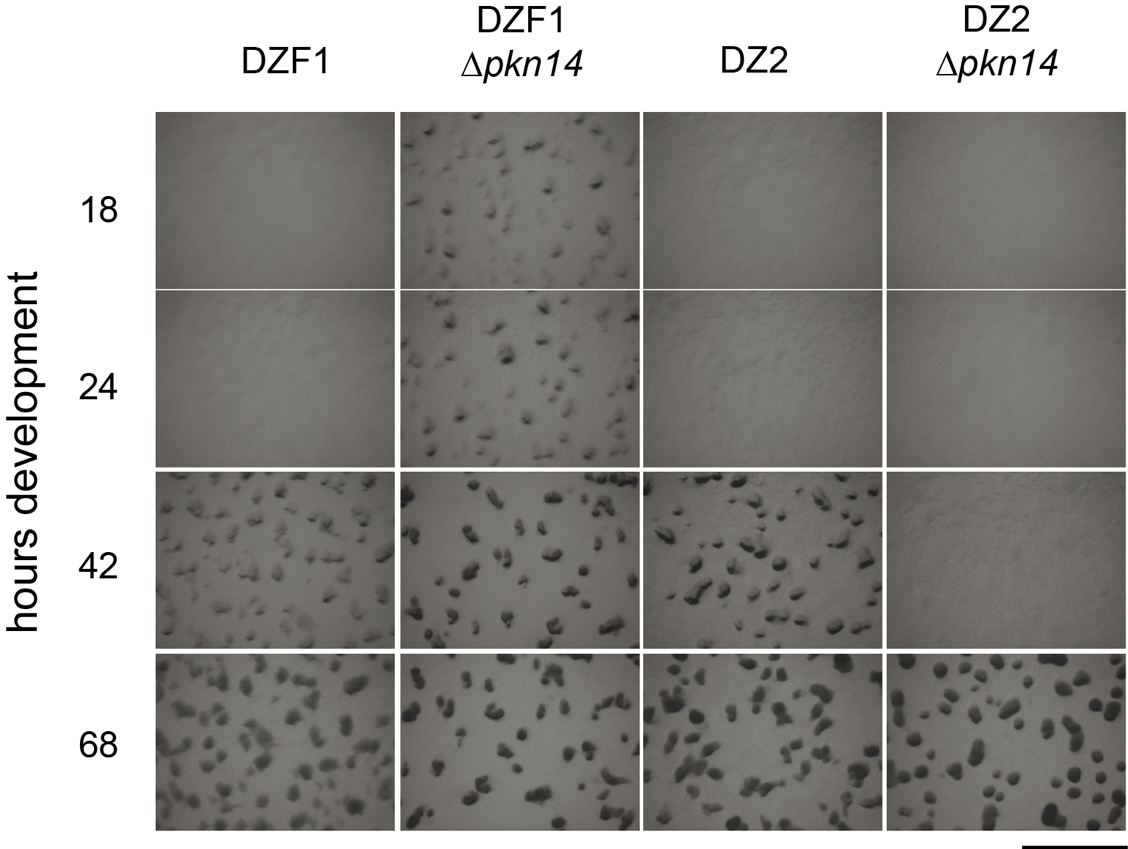

Figure S3. Developmental phenotype of Δ*pkn14* in the DZF1 versus DZ2 backgrounds. Generation of the *pkn14* deletion in the DZF1 background recapitulates the early development phenotype previously observed by Nariya et al. (Nariya & Inouye, 2005b). 4 x 10^8^ cells of DZF1, DZF1 Δ*pkn14* (PH1550), DZ2, or DZ2 Δ*pkn14* (PH1132) strains were spotted on CF agar plates (Campos & Zusman, 1975), and images were recorded at the indicated hours of development at 32 ^ο^C. Bar: 1.0 mm.

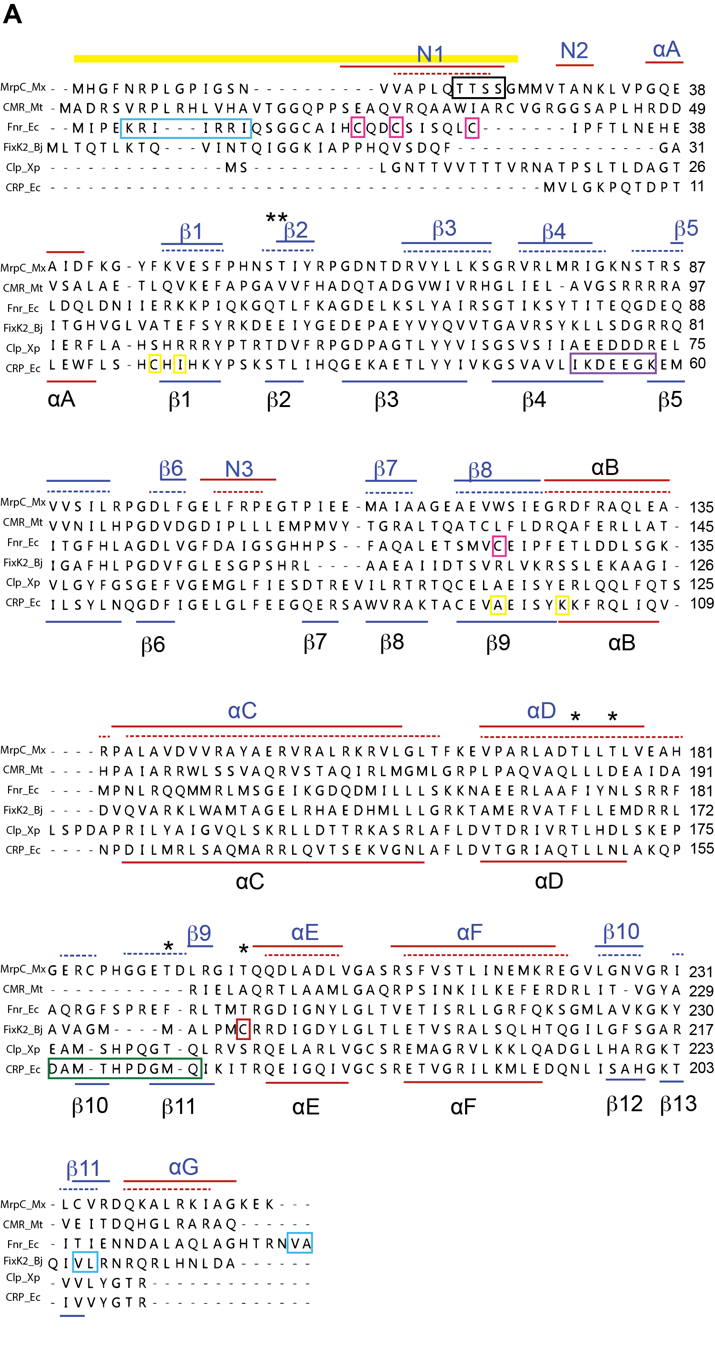

Figure S4. Alignments of MrpC with Crp/Fnr homologs reveals functional residue conservation. A. Clustal Omega (Sievers *et al.*, 2011) alignments of *M. xanthus* MrpC (MrpC_Mx), *Mycobacterium tuberculosis* Cmr (CMR_Mt), *E. coli* Fnr (Fnr_Ec), *Bradyrhizobium japonicum* FixK_2_ (FixK2_Bj), *Xanthomonas campestris* (Clp_Xp), and *E. coli* Crp (CRP_Ec) were analyzed in MegAlign Pro 14.0.0 (DNASTAR) with 1000 iterations of refinement using algorithm-based HMMs. Secondary structure helices (red lines) and beta-sheets (blue lines) from the crystal structures of *E. coli* Crp and *M. tuberculosis* Cmr crystal structures are shown below and above the alignments, respectively. Secondary structures predicted by Phrye2 intensive homology modeling and depicted by QtMG are shown by dashed lines above the alignments for M. xanthus MrpC. Thick yellow line: *M. xanthus* MrpC N25; black box: *M. xanthus* MrpC TTSS motif; blue boxes: residues necessary for proteolytic targeting; pink boxes: Fnr residues coordinating the Fe-S cluster; red Box: FixK_2_ cysteine residue oxidized to control DNA binding; green box; *E. coli* CRP activating region 1; yellow boxes: *E. coli* CRP activating region 2; purple boxes: *E. coli* CRP activating region 3; asterisks: Pkn14-dpendent phosphorylated residues identified by mass spectrometry analysis on *M. xanthus* MrpC.

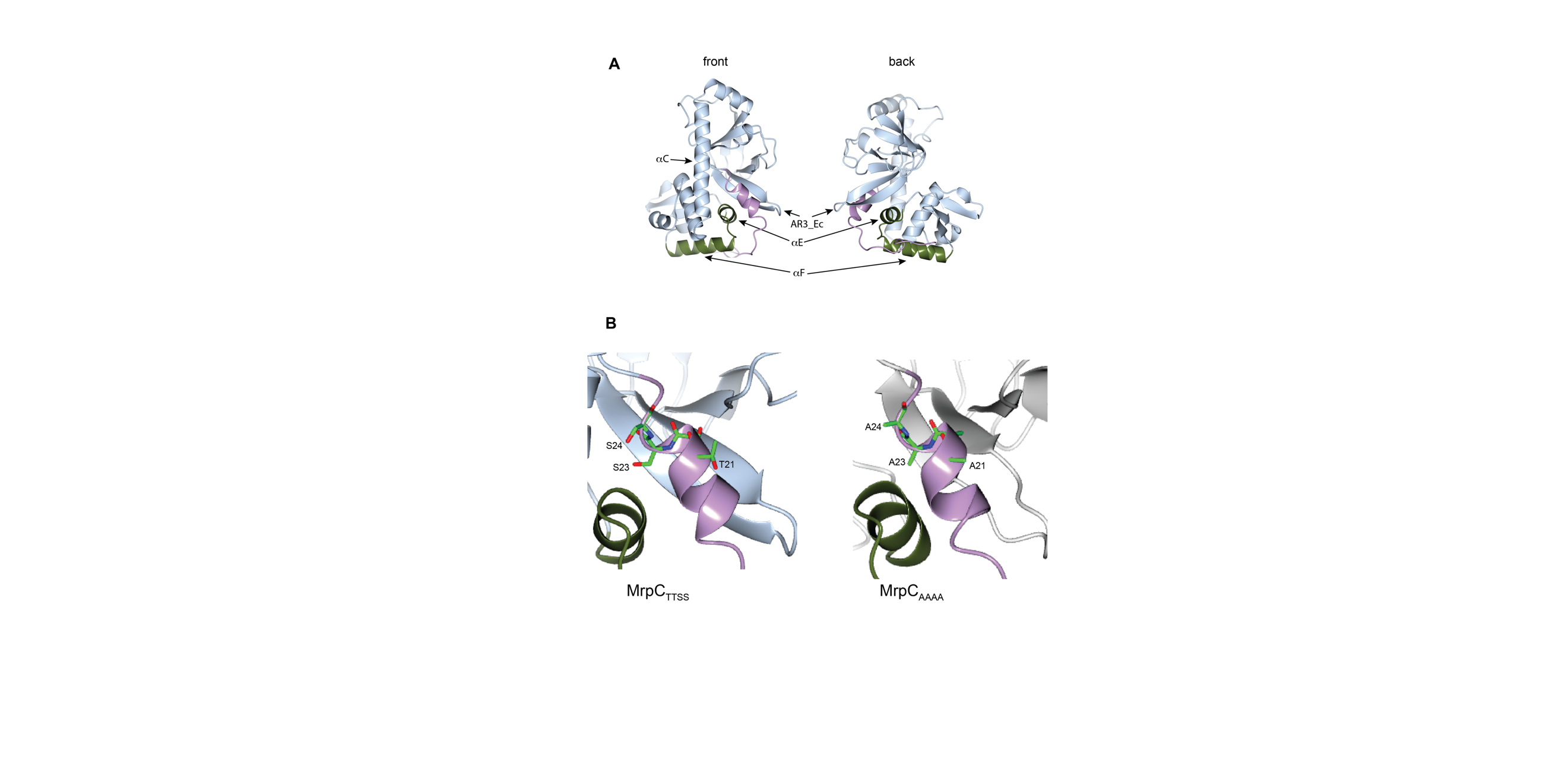

Figure S5. Predicted MrpC tertiary structure. A. Phyre2 (Kelley *et al.*, 2015) intensive mode was used to model MrpC. 94% of the protein was modeled at >90% confidence; residues 1-15 were modeled ab initio and considered unreliable. Structures were viewed and colored in CCP4MG version 2.10.10 (McNicholas *et al.*, 2011). The DNA binding helix-turn-helix (E and F helices) are depicted in green. The amino terminal residues 1-25 are depicted in purple. Note the amino terminal extension (N25) threads between the dimerization (C helix) and the beta scaffold with the TTSS motif at the end of a helix. This region is similar to the *M. tuberculosis* Cmr N1 helix (Protein Data Bank accession code 5W5A)(Ranganathan *et al.*, 2018). B. Close view of residues of the TTSS motif in wild type MrpC (MrpC_TTSS_) (left) or the MrpC_AAAA_ mutant (right). The respective residues 21-24 are rendered as sticks with O, N, and C atoms depicted in red, blue, and green, respectively.
